# Spatial propagation of temperate phages within and among biofilms

**DOI:** 10.1101/2023.12.20.571119

**Authors:** James B. Winans, Sofia L. Garcia, Lanying Zeng, Carey D. Nadell

## Abstract

Bacteria form groups comprised of cells and a secreted polymeric matrix that controls their spatial organization. These groups – termed biofilms – can act as refuges from environmental disturbances and from biotic threats, including phages. Despite the ubiquity of temperate phages and bacterial biofilms, live propagation of temperate phages within biofilms has never been characterized on cellular spatial scales. Here, we leverage several approaches to track temperate phages and distinguish between lytic and lysogenic host infections. We determine that lysogeny within *E. coli* biofilms initially occurs within a predictable region of cell group packing architecture on the biofilm periphery. Because lysogens are generally found on the periphery of large cell groups, where lytic viral infections also reduce local biofilm cell packing density, lysogens are predisposed to disperse into the passing liquid and are over-represented in biofilms formed from the dispersal pool of the original biofilm-phage system. Comparing our results with those for virulent phages reveals that temperate phages have previously unknown advantages in propagating over long spatial and time scales within and among bacterial biofilms.

## Introduction

Across large ranges in spatial scale and phylogenetic distance, many species - from microbes to the largest metazoans - form groups as an adaptive strategy to navigate environmental challenges^1–5^. Microbial collectives, often termed biofilms, are produced either in a free-floating state or attached to surfaces. Biofilms are primarily composed of cells and a suite of secreted polymers, called the extracellular matrix^6–8^. This mode of group formation among bacteria is ubiquitous in both natural and human-made environments, and a primary function of collective growth is protection against invading competitors, diffusible antimicrobial compounds, predatory bacteria, and phages^9–11^. The microscopic scale dynamics of group protection against these threats are not well known, particularly for phages and predatory bacteria^12–15^. Complementing prior work with microscopic observation of phage-biofilm interactions is essential to provide insight into the driving mechanisms of these interactions at the spatial scale on which microbial phenotypes directly manifest.

Microbial predators have a broad diversity of life history patterns. Obligate lytic phages, also called virulent phages, inject their genome into host cells, where the cellular machinery is coopted for phage replication preceding host cell lysis^16^. Temperate phages, by contrast, have two different modes of propagation: 1) they may proceed with a lytic infection like virulent phages, killing the host cell and producing a new burst of phage virions, or 2) they may integrate into the bacterial genome or persist as an episome to generate a host lysogen that carries the phage’s genome along with it as it undergoes normal growth and division^17^. Once integrated into a host’s genome, prophages can introduce new genes that substantially alter cell physiology, including antibiotic resistance, metabolic capacity, and virulence^18–21^. Additionally, the movement of temperate phages into and out of host genomes often mediates horizontal gene transfer^22^; has powerful, ancient, and widespread effects on microbial evolution^23,24^; and can drive lysogenic conversion of pathogens that present major challenges to human health^25^. An enormous wealth of knowledge of phage ecology, evolution, and molecular genetics has been gathered over a century of research^16,26–30^. However, despite the ubiquity of biofilms and phages in microbial ecology, we are still learning the fundamentals of how phages interact with biofilm-dwelling hosts. Temperate phage propagation, in particular, remains uncharacterized at cellular and multicellular spatial scales among host bacteria in biofilms.

Recent work has begun to clarify the fundamentals of lytic phage propagation at high spatial and temporal resolution in the biofilm context. This research has shown that, for several species, bacteria within biofilms are often protected from lytic phage exposure^31–35^, and that this protection is dependent on biofilm growth and architectural maturity at the time of phage introduction. When *E. coli* is growing in a biofilm context, for example, the hosts’ interactions with lytic phages are profoundly different relative to shaken liquid culture conditions^12,31,36^. Mature *E. coli* biofilms can maintain net-positive growth in the presence of T7 phages by blocking phage diffusion in a manner dependent on the secretion of curli amyloid fibers^31^, a core proteinaceous component of the *E. coli* biofilm matrix. Phages can be trapped within the networks of highly packed cells and curli fiber protein on the periphery of the biofilm, but the phages remain viable and can infect susceptible bacteria attempting to colonize the biofilm exterior from the planktonic phase^36^. The spatial constraint on phage and cell movement within biofilms inherently tends to create negative frequency-dependent selection for genetic phage resistance – that is, resistant mutants are selectively favored when rare, but disfavored when common. This result occurs because genetically resistant cells, once they are common, further reduce phage mobility in the system and can prevent susceptible cells from being exposed and infected^37,38^.

The work summarized above has established an early picture of how obligate lytic phages interact with biofilm-dwelling hosts at microscopic spatial scales. In contrast, there is very little analogous work on how temperate phages propagate through biofilm communities, nor has prior research addressed how the patterns of phage propagation at the community scale are driven by cellular scale interaction dynamics. Some previous research has used traditional macroscopic measurements to explore how temperate phage exposure influences biofilms^39^. A recent study showed that non-specific genome insertion by prophages can accelerate bacterial evolution by increasing host mutation rate in biofilms of *P. aeruginosa*^40^. Other work with lambdoid phage H-19B and *E. coli* MG1655 indicated that almost all host cells are quickly lysogenized in biofilm populations^41^. This work set a precedent that temperate phage exposure in biofilms can be important; but efforts thus far have used techniques that disrupt the physical architecture of the biofilms and average over the entire host and phage populations. Crucial questions therefore remain: how do temperate phages propagate through biofilms at the cellular scale; how do these fine-scale patterns influence population and community structure and composition; and how are the answers to these questions distinct for temperate phages relative to obligate lytic phages^22,42–44?^

To begin answering these questions, we developed a model system in which host bacteria can be visualized at single cell resolution, and in which naïve (uninfected) cells, lysogenized cells, ongoing lytic infections, and phage virions can be distinguished by fluorescence microscopy. The model comprises monoculture biofilms of *E. coli* AR3110 cultivated in microfluidic devices under constant media flow, to which temperate phage λ (or, in selected control experiments, obligate lytic phage T7 or λΔ*cI*) can be added at any time during or after host cell surface colonization, cell growth, and biofilm matrix secretion. *E. coli* is among the most extensively studied organisms in microbiology, and its mechanisms of biofilm formation have likewise been dissected thoroughly^45–52^. Similarly, temperate phage λ has been studied in detail over many decades of work^53–63^. Here, we assess how this phage and its host interact within biofilms at the resolution of individual cells and phage virions. Our study yields new insight into the principles of temperate phage propagation at cellular and multicellular biofilm scales, the impact of the temperate phage life history strategy on host biofilm architecture, and the nature of local versus distal dispersal-based spread of temperate phages versus virulent phages within and among biofilms.

## Results

*E. coli* AR3110 was engineered to constitutively produce the far-red fluorescent protein mKate2. AR3110 is a K-12 derivative with biofilm matrix production restored relative to the parental W3110 K-12 lineage^45^. The resulting strain was used to inoculate microfluidic devices composed of glass coverslips bonded to polydimethylsiloxane molds; chambers measured 5000 μm long, 500 μm wide, and 70 μm in height from glass substratum to PDMS ceiling **(Supplementary Fig. 1)**. After a 45-min period of stationary conditions to allow cells to attach to the glass, M9 minimal media with 0.5% maltose was introduced at a flow rate of 0.1 μL min^-1^ (average flow velocity = 45 μm/s) for 48 h. We chose maltose as the sole carbon source to ensure that cells produce the LamB maltoporin, which is the attachment site for phage λ^64^. The media influent was then switched to a new syringe with the same growth media plus λ phages at a concentration of 10^4^ PFU μL^-1^ for 24 h. This strain of λ, λLZ1367, has a copy of the teal fluorescent protein mTurquoise2 translationally fused to λD, which encodes the capsid protein gpD^65,66^. λLZ1367 also contains a downstream transcriptional fusion of *mKO2* to the *cI* locus encoding Repressor, which causes lysogenized hosts to produce the orange fluorescent protein mKO2^66–68^. Overall, this arrangement of constitutive and regulated fluorescent protein constructs on the bacterial and phage genomes allows for high resolution spatial tracking of non-lysogenized hosts (mKate2), lysogenized hosts (mKO2), lytic hosts and λ phage virions (mTurquoise2). By comparing total fluorescence of serial dilutions of mTurquoise2-labeled phage virions to PFU titers, we confirmed that the teal fluorescence intensity of λLZ1367 scales linearly with phage titer in our image data (**Supplementary Fig. 2)**. Furthermore, the production of mKO2 allows us to accurately measure the fraction of lysogenized cells for all experiments below, which we confirmed by comparing lysogen quantification via mKO2 fluorescence thresholding to measurements of lysogen CFU counts within control chambers (**Supplementary Fig. 2**).

### Lytic infections and host lysogenization are restricted to the biofilm periphery in a manner dependent on biofilm matrix and cell packing

As biofilms increase in size under flow, cells may detach actively or passively from existing groups, re-attach elsewhere in the chamber, and begin producing new cell clusters. As a result, after 48 h of incubation prior to adding phages, our biofilm chambers contained a variety of *E. coli* host cell groups of different size and packing architecture (**Supplementary Fig. 3**). In this culture condition, the largest and longest-established clusters of *E. coli* are each composed of a densely packed center, surrounded by cells with lower packing density (**Supplementary Fig. 3**). After phages were added to the flow devices, spatial heterogeneity in cell infection state (unexposed, lysed, lysogenized) arose within 24 h. Over this time scale, per expectation, the predominant mode of phage propagation was through lytic infection events in which phages produced a new burst of phage virions. Consistent with previous reports, we found that biofilm cell groups of relatively small size and low cell packing were lysed and removed from the system^12–14,31^. Larger, more densely packed clusters of *E. coli* were commonly found to have a peripheral region containing many λ phages, newly formed lysogens, and naïve host cells, as well as a highly packed central core of cells with few phages and no lysogenized hosts (**Fig. 1A, C**). Previous work has shown that the *E. coli* matrix protein curli is essential for the cell-cell packing that can slow or halt phage diffusion^31^. We reasoned that if curli matrix production is necessary for creating biofilm regions that are inaccessible to phages, a mutant strain of *E. coli* lacking the ability to produce curli should produce biofilms in which all cells are phage-accessible, and lysogens should arise relatively homogeneously throughout the system. We tested this possibility by repeating the experiments above using a strain of *E. coli* (denoted Δ*csgBA*) harboring deletions of *csgB* (encoding the curli baseplate) and *csgA* (encoding the curli amyloid monomer). This strain does not have a growth defect in liquid culture; however, its biofilms do not grow to the same height as those of WT, and their maximal cell packing is lower than that of the WT strain; these differences occur because Δ*csgBA* cells are not as well attached to each other and the underlying glass surface as WT curli-producing cells^69^. When we repeated our experiment with biofilms produced by the Δ*csgBA* double mutant – consistent with our hypothesis and prior literature – phages and lysogens appeared to be homogeneously distributed among host cells (**Fig. 1B**).

**Figure 1.**
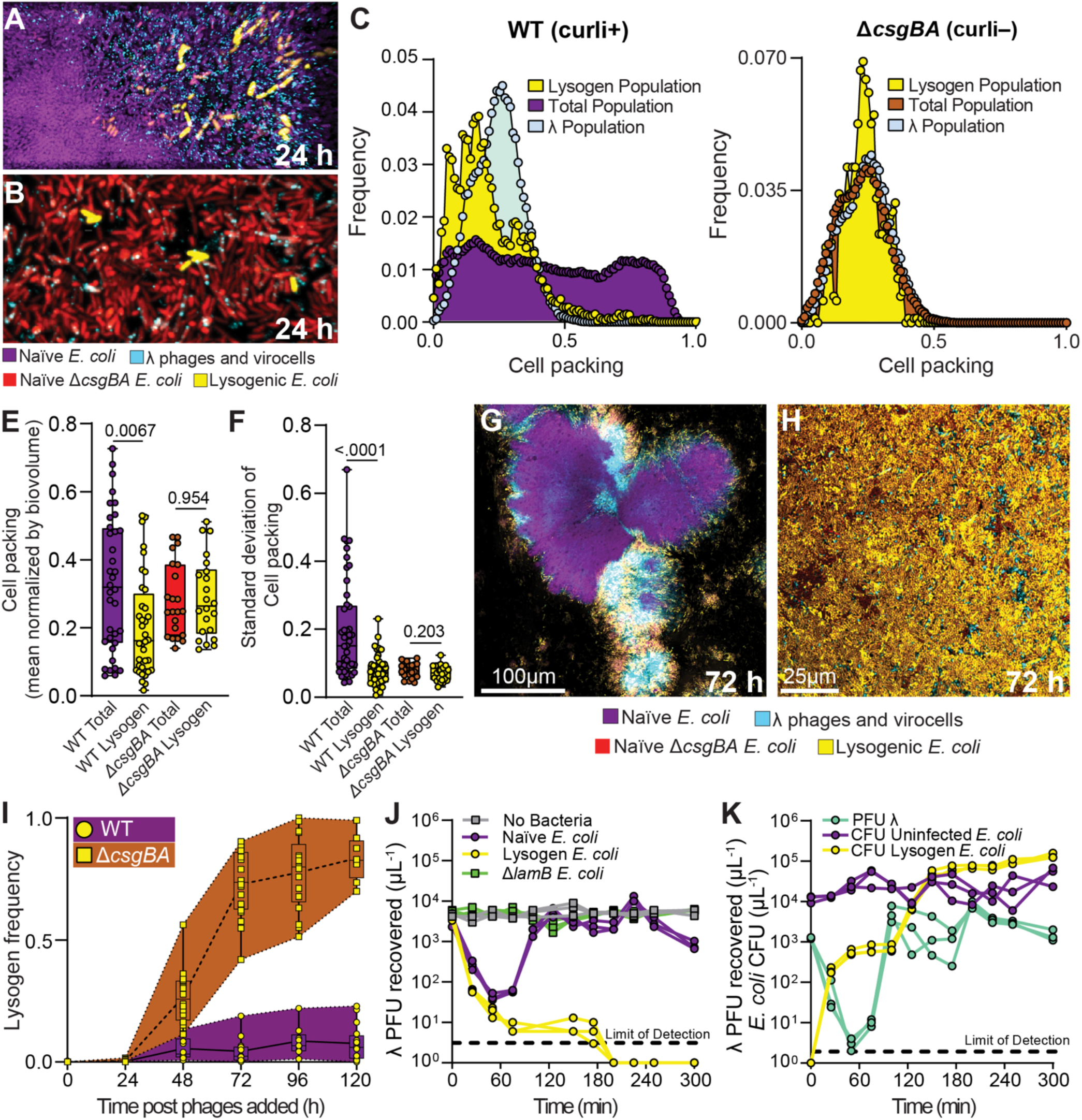
(A) WT *E. coli* biofilms (purple) invaded with λ phages (turquoise) for 24 h, with some cells becoming lysogenized (yellow). (B) Δ*csgBA* biofilms (red) invaded with λ phages (turquoise) for 24 h, with some cells becoming lysogenized (yellow). Images in (A) and (B) are 3-dimentionsal renderings (for WT: 64Lx32Wx30H μm; for Δ*csgBA*: 64Lx32Wx8H μm). (C,D) The frequency distributions of phages, lysogens, and total host cell populations with respect to biofilm cell packing for (C) the WT biofilm experiments and (D) the Δ*csgBA* biofilm experiments. (E,F) Quantitative comparisons of the mean cell packing and the standard deviation in cell packing near lysogenized cells in comparison with the total host bacterial population over replicate runs of the experiments shown in (A-D). (G, H) Representative 2-dimensional optical section images of experiments in which biofilms were treated as for (A,B) but were tracked for 120 h instead of 24 h. (I) Lysogen frequency over 120 h for the WT background and the Δ*csgBA* background. (J) Phage titer over 300 min when inoculated with no bacteria, naïve *E. coli*, lysogenized *E. coli* (phage resistant and phage-adsorbing), and Δ*lamB E. coli* (phage-resistant, but not phage-adsorbing). (K) Bacterial CFU count for naïve host *E. coli* and lysogenized *E. coli*, as well as λ phage titer over 300 min when phages are introduced to a population of only naïve hosts in shaken liquid culture.

To determine whether the naïve *E. coli* cells that survived did so due to *de novo* evolution of resistance to phage λ, we repeated the experiment above, but after introducing λ phages for 24 h, all of the cells within the microfluidic device were flushed out and plated for resistance to λ infection (See Methods). The frequency of λ resistance in this surviving *E. coli* population was approximately 10^-5^, which is not significantly different from the background frequency of resistant mutants in our overnight cultures used to inoculate the biofilm chambers at the start of the experiment (**Supplementary Fig. 4**). This result indicates that *de novo* genetic λ resistance did not contribute to the population dynamics in our experiments, and it is consistent with the fact that our biofilm cultures contain bacterial and phage population sizes and contact rates that – due to spatial constraint on bacterial and phage movement – are orders of magnitude lower than those found in shaken liquid batch culture^12,31,36^. The results above instead suggest – consistent with our prior work exploring T7 phage propagation – that *E. coli* biofilm architecture can limit the diffusion of phages to a region along the periphery of cell groups and create a refuge against phage exposure in its core region^12,31^. The biofilm cell packing architecture that leads to phage protection only occurs if *E. coli* has sufficient time to grow prior to the introduction of phages; if phages are introduced from the beginning of biofilm growth, the spatial pattern of naïve cell distribution, cell packing, phage diffusion, and lysogen localization shown in Figure 1 does not occur (**Supplementary Fig. 5**).

Our observations above suggested that for *E. coli* biofilms of sufficient size and architectural maturity, temperate phage propagation can occur along the peripheral regions of cell clusters, where cell packing is lowest. In general, cell packing increases with depth from the biofilms’ outer surface (**Supplementary Fig. 6**), but we will use cell packing as the key index for this study, as cell packing, rather than biofilm depth, is more directly mechanistically linked to phage localization. The lytic activity of the phages tends to reduce local cell packing even further, and it is within this region of reduced cell density that lysogenization is observed (**Supplementary Fig. 7**). To assess this visual intuition quantitively, we used the BiofilmQ analysis framework to calculate the distribution of lysogenized cells with respect to local cell packing density and to compare their distribution with that of the total *E. coli* population (See Methods, **Supplementary Fig. 8**). We found that lysogens are indeed over-represented in biofilm regions with lower packing density and are heavily skewed in this respect compared to the background total distribution of cell packing within large *E. coli* biofilm colonies (**Fig. 1 C**). Interestingly, the frequency distribution of lysogens peaks at lower biofilm cell packing than the λ phages themselves, which indicates that lysogenization has yet to equilibrate with the spatial range of phage spread by 24 h (see below for longer time course experiments). When we repeat this experiment with a Δ*csgBA* strain lacking curli production, the spatial distributions of phages and lysogens relative to local cell packing are indistinguishable from that of the entire host cell population (**Fig. 1D, Supplementary Fig. 8**). The spatial patterns of lysogen and phage localization for WT versus Δ*csgBA* host background were consistent across many runs of these experiments (**Fig. 1E, F**). Notably, we observed lysogens within Δ*csgBA* biofilms at a marginally higher average cell packing than we did for lysogens within WT biofilms. We speculate that this was due to the ability of WT biofilms to progressively halt phage diffusion as they approach regions of high cell packing; Δ*csgBA* biofilms, on the other hand, pose no or minimal obstacles to phage diffusion and allow them to enter their full volume and range of cell packing density (**Supplementary Fig. 9**).

The results thus far suggested to us that the spatial patterning of lysogenization was due to the cell packing architecture that *E. coli* produces, which blocks phage diffusion into the highly packed core regions of host biofilm colonies. Previous research documenting halted diffusion of T7 phages into *E. coli* biofilms showed that this protective phage blocking was a function of the packing structure generated by the secretion of curli matrix proteins^31^. A difference we observe here is that *E. coli* biofilm cell groups become larger with rougher surface topography in our media conditions (M9 minimal with 0.5% maltose), as opposed to the flat mat architecture seen in the prior study of T7 diffusion in *E. coli* biofilms (which was done in 1% tryptone broth). To document curli production quantitatively in our culture conditions, we used a previously engineered strain of *E. coli* with a fluorescent transcriptional reporter inserted into the *csgBAC* operon^31^. We found that *csgBAC* transcription increases with cell packing up until the very highest values of cell packing fraction that we measured (**Supplementary Fig. 10**)^70,71^. We repeated these experiments with an *E. coli* strain that has a 6xHIS tag fused to the csgA monomer, which allows for immunostaining of the curli biofilm matrix. This epitope tag has been shown not to interfere with the functionality of the curli matrix^31^. We observe a similar relationship between cell density and curli immunostaining as we did with the *csgBAC* transcriptional reporter (**Supplementary Fig. 10**). In addition to documenting the production of curli matrix by *E. coli* in our culture conditions, we note that high *csgBAC* transcriptional activity at the tightly packed center of biofilms in our experiments suggests that cells on the interior remain physiologically active and are not avoiding phage infection due to nutrient starvation and quiescence. Previous work has also documented that, at the center of biofilms of the size order that we examine here, cell still have access to growth limiting substrates and remain physiologically active^12,72^.

### Lysogen abundance and localization equilibrate in biofilms over multiple days

Our experiments thus far suggest that by virtue of matrix secretion and cell packing architecture in established *E. coli* biofilms, the introduction of temperate phage λ leads to lytic proliferation in a finite peripheral region of sufficiently large, densely packed biofilm cell groups. An inevitable byproduct of this pattern is that lysogenized hosts tend to occur in the outer boundary regions of large biofilm clusters, where cell-cell packing is lower by default and is decreased further by lytic-cycle phage activity (**Supplementary Fig. 7**). We were next curious about the stability of these patterns through time.

We grew WT and *ΔcsgBA E. coli* biofilms for 48 h prior to introducing λ phages continuously for an additional 120 h, acquiring images of replicate biofilm chambers daily (**Fig. 1G, H**). The frequency of lysogens within WT biofilms increased gradually before reaching a steady state at ∼10% of the population after 96 h (**Fig. 1I**). Given the constant influx of phages into the system, and the fact that lysogens themselves grow and divide in the peripheral regions of biofilm with highest nutrient availability, it was not immediately clear why lysogenized cells would tend to equilibrate at 10% of the population rather than continuing to increase in relative abundance. The first and obvious factor limiting lysogenization – explored in Figure 1 – is that lysogen frequency can reach an upper limit within structurally mature biofilms due to phage diffusion constraints that limit what fraction of the original cell population is phage-exposed. As a result, uninfected cells within a cell group can replicate and keep pace with the growth of newly replicating λ lysogens around the cell group periphery. A second explanation, not mutually exclusive from the first, is that newly formed lysogens surrounding uninfected cells could create barriers against further phage propagation, as lysogenized cells can adsorb λ virions but are immune to superinfection^73,74^. An additional explanatory factor, also not mutually exclusive from the previous two, is that newly formed lysogens might be disproportionately pre-disposed to disperse from biofilms due to their spatial arrangement on the shear-exposed biofilm periphery. We assess the first two possibilities below and expand upon the third in subsequent sections.

To first assess the role of *E. coli* biofilm structure on lysogen proliferation over time, biofilms of Δ*csgBA E. coli* were grown for 48 h without phages, and then monitored for an additional 120 h under continuous λ exposure. As observed for the WT biofilms above, lysogens initially arise slowly and after 48 h comprise only a small fraction of the total population. By contrast, after 48 h of the phage exposure the relative abundance of lysogens within Δ*csgBA* host biofilms continues to increase sharply, departing more and more dramatically from the pattern seen for WT host biofilm structure. By 96 h after introduction of phages, lysogens constitute 80-100% of the total Δ*csgBA* population (**Fig. 1I**). This result makes it clear that matrix structure is a central factor controlling phage movement and replication, but it does not exclude the possibility that lysogens on the periphery of biofilms reduce further phage entry by adsorbing them.

To assess the adsorption of free λ phages to different host *E. coli*, we turned to well-mixed batch culture experiments in which λ phages were incubated with λ-susceptible naïve *E. coli*, λ-lysogenized *E. coli* (which can adsorb phages but do not support lytic propagation), Δ*lamB E. coli* (which lack the LamB maltose importer to which phage λ binds, and therefore do not adsorb phage λ), or a blank media control. We began these experiments with an initial phage:bacteria ratio of 1:10 (multiplicity of infection = 0.1) and tracked phage adsorption and amplification by quantifying free λ phage titer from the liquid media in each culture condition every 25 min for 5 h (see Methods). As expected, incubating phage λ in sterile media or with the Δ*lamB E. coli* deletion mutant produced no change in phage titer throughout the experiment. When we incubated λ with a previously lysogenized *E. coli* population, we observed a rapid decline in liquid phage titer that fell below the limit of detection by 200 min; this follows expectation and recapitulates the classic superinfection immunity result^74–76^. When we repeated this experiment with λ-susceptible *E. coli*, we observed an initial drop in phage titer as phages adsorbed to host cells. However, in this condition, the initial drop in phage titer is followed by a >100-fold increase corresponding to a burst of new phages from a round of lytic infection. As explored below, a rapid increase in new *E. coli* lysogens also occurs at this time **(Fig. 1J)**. From this time point forward the phage-host population dynamics become more complex, as the host *E. coli* population comprises naïve host cells and newly lysogenized hosts (both of which are actively growing), in addition to free λ phages. To better understand these dynamics, we repeated this experiment and quantified the changing abundances of lysogenized hosts and non-lysogenized naïve hosts by CFU count, in addition to monitoring free λ phage titer (see Methods). Lysogen abundance increases sharply straight from the start of the experiment, saturating until the release of new phages from the first round of lytic infection bursts from the initially naïve *E. coli* host cell population. With the release of this new λ phage burst, lysogen abundance again increases, due to growth of existing lysogens and new lysogenization of naïve cell hosts. Surprisingly, the non-lysogenized, uninfected naïve *E. coli* cells showed variance of less than 1 order of magnitude from their inoculated population size. Our interpretation of this result is that because the naïve cells are actively replicating and partially being converted to lysogens, which adsorb free phages, their populations remain relative stable despite successive rounds of phage release, adsorption, and lysogenization or lytic amplification (**Fig. 1K, Supplementary Fig. 11**)

Taken together, these results suggested that the steady state frequency of lysogens within naïve biofilms exposed to phages can be explained by phage diffusion limitation conferred by matrix secretion and cell packing structure, in addition to a smaller contribution from phage shielding by lysogens that adsorb free λ virions. This interpretation predicts that in biofilms from which high cell-packing architecture and the ability of lysogens to adsorb phages have been removed, phages should be able to access all infectible cells by diffusion. We tested this idea by repeating the biofilm experiment from this section with a slightly modified protocol. We cocultured Δ*csgBA* cells and a Δ*csgBA* Δ*lamB* mutant to mimic the condition in which Δ*csgBA* naïve cells grow alongside a phage-immune strain, but in this case the Δ*csgBA* Δ*lamB* phage-immune strain cannot impede phage movement by adsorption. We found that after 24 h of phage inoculation, the Δ*csgBA* cells had significantly higher overlap with phages compared to the Δ*csgBA* Δ*lamB* cells (**Supplementary Fig. 12**). When we performed this same experiment with Δ*csgBA* and a Δ*csgBA* lysogenized with λ (which is phage resistant and can absorb/neutralize free λ virions), we found no difference in phage overlap between the two strains (**Supplementary Fig. 12**). The comparison between these two experiments indicates to us that phage adsorption by phage-immune cells can reduce phage mobility and contribute to maintaining a residual uninfected population of susceptible cells.

### Lysogenized cells are more likely to disperse due to their characteristic spatial arrangement

So far, we have explored the process of phage propagation within individual biofilm patches as a function of the local cell packing architecture and the adsorption of phages by either naïve or lysogenized host cells as the biofilms grow toward steady state. We were also curious as to the flux of dispersed lysogens from biofilms’ exterior, particularly in light of the finding that lysogens tend to be created along the biofilm periphery in naïve biofilms exposed to phages. Given that lysogens are skewed in their spatial distribution toward peripheral regions of lower cell packing, and that ongoing lytic cycle activity by phage λ reduces local cell packing further in the regions where lysogens are located (**Supplementary Fig. 7**), we hypothesized that lysogens are disproportionately more likely to disperse from cell groups and re-colonize elsewhere relative to their total frequency in biofilms.

To test this idea experimentally, we incubated WT *E. coli* biofilms as in Figure 1, inoculated them with phages for 24 h, and imaged the system to measure abundance of naïve and lysogenized cells (**Fig. 2A**). We then immediately collected effluent from these microfluidic devices and prepared the contents for imaging with no delay during which cell growth or changes in phage activity could occur. We calculated the relative abundance of lysogens in the liquid effluent and found that indeed the lysogen frequency in the exiting liquid phase is significantly higher than it is in the total biofilm population within the chambers (**Fig. 2B,C, Supplementary Fig. 13**). Specifically, lysogens were 10-fold more likely to be dispersed than the average non-lysogenized naïve host cell in the system. On the other hand, if we isolate only a fraction of the population residing at a cell packing fraction of less than 0.3, then the lysogens’ frequency in this subpopulation becomes indistinguishable from its frequency in the effluent (**Fig 2C**). This observation reinforces the interpretation that the dispersing cells are derived primarily from the outermost, lower-density biofilm regions, which is where the lysogens are most abundant. Direct quantification confirms that phages and lysogens are indeed restricted to the outer layers of biofilms over many replicates of this experiment (**Figure 2D,E, Supplementary Fig. 6**). If we conducted this same experiment after 72 h of phage exposure (as opposed to 24 h from the experiments above), the disparity between lysogen frequency in the effluent relative to that in the original biofilm was even larger (**Supplementary Fig. 14**). This suggests that the pattern of biased dispersal of lysogens is a persistent feature of this experimental system, which strengthens over time as more lysogens are produced by phage-host contact events and division of existing lysogens along the biofilm periphery.

**Figure 2.**
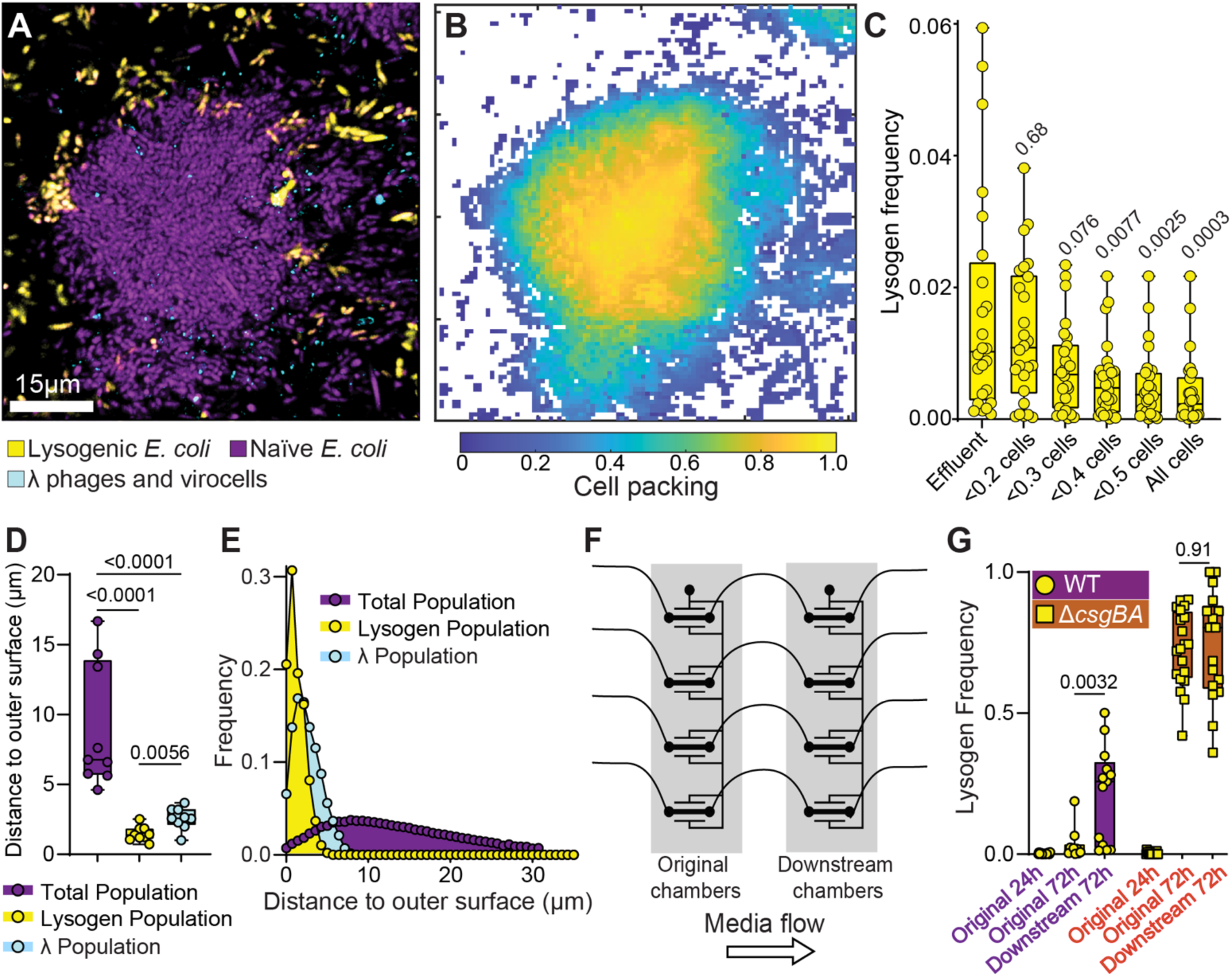
(A) Representative 2-dimensional optical section of a phage-naïve WT *E. coli* (purple) biofilms following exposure to phages (cyan). Lysogens (yellow) have been formed around the periphery of the biofilm cell group. (B) A heatmap showing a 2-dimentional projection of cell packing measurements for the full 3-dimensional z-stack of images capturing the group of cells shown in (A). (C) A comparison of the frequency of lysogens in liquid effluent exiting biofilm chambers, in comparison with the frequency of lysogens in regions of different cell packing. Effluent lysogen frequency is closest to that in the regions of lowest cell packing on the periphery of biofilm clusters. (D) Measurements of distance to outer cell cluster surface for the total prey cell population, lysogens, and phages. (E) Frequency distributions of distance to outer cell group surface for the total prey population, lysogens, and phages across all experimental replicates. (F) An illustration of the dispersal assay in which biofilms are first grown in the left hand (‘original’) chamber, after which the effluent from this chamber is used to inoculate a second (‘downstream’) chamber to simulate dispersal from one location to another. (G) The frequency of lysogens in the original upstream chamber and the new downstream chamber for dispersal experiments with the WT *E. coli* strain background and the isogenic Δ*csgBA* mutant that cannot produce curli matrix.

As phage λ carries genes that can modify host outer membrane composition^77,78^, we checked whether λ lysogens are more likely to disperse due to a property conferred by the integrated phage genome itself. To do so we inoculated biofilm chambers with a 1:1 mixture of pre-lysogenized cells and naïve cells and tracked the biovolume of both strains within the chamber and within the effluent. We found that uninfected and lysogen cells compete neutrally when in biofilm culture together (**Supplementary Fig. 14**); that is, net results of population growth, minus total cell deaths, minus cell dispersal events are the same for WT and lysogens when they are inoculated together at the start of the experiment. These data imply that in the case when phages are added to an initially phage-naïve, non-lysogenized host biofilm, lysogens disperse more frequently than naïve cells not because of a change in biofilm dispersal physiology caused by prophages, but rather due to the spatial arrangement in which lysogens are generated. This interpretation predicts that in Δ*csgBA E. coli* biofilms, which lack the cell-cell packing structure that creates stratified lysogen distributions (Figure 1), there should be no discrepancy between within-biofilm lysogen abundance and liquid effluent lysogen abundance. This prediction was confirmed in control experiments **(Supplementary Fig. 14)**.

The result that lysogens are especially disposed to disperse from naïve WT biofilms prompted us to consider larger spatial scale patterns of phage spread and biofilm distribution. We hypothesized that because lysogens are disproportionately more likely to depart from biofilms relative to non-lysogenized cells, the new biofilms re-seeded by this dispersal pool downstream from the original population will be over-represented for lysogens relative to biofilms from which the dispersed cells originated. We tested this possibility by growing biofilms as above for 48 h, adding phages continuously for 24 h, sampling effluent from these chambers, and inoculating the liquid effluent into fresh microfluidic chambers (**Fig. 2F**) . This effluent contained mixtures of naïve cells, lysogenized cells, and free λ phages. After a 45 min attachment phase, we incubated these biofilms for 72 h without any additional phages being introduced to the flow devices. After 72 h of growth, the frequency of lysogens in the downstream chambers was significantly higher than what we observed in the original chambers (**Fig. 2G**). There was notably high variance in this result; we speculate that this is due to biological variation in the upstream biofilm population structure and the bottleneck re-colonization events from the effluent samples, which deposited relatively small, noise-prone samples of cells on the glass in downstream biofilm chambers. Variation in the number of phages departing the original chambers and interacting with populations of bacteria on their way to the new chambers could have contributed to variation in downstream lysogen abundance as well. When we performed this same experiment with our Δ*csgBA* strain of *E. coli*, we saw no difference between the original chambers and downstream chambers at equivalent time points, again consistent with all of the analyses above (**Fig 2G**).

Altogether, our data suggest that the natural spatial constraints of biofilm cell group architecture lead to a repeatable spatial pattern of lysogen generation when naïve biofilms are exposed to temperate phages. Lysogens tend to be concentrated in the periphery of established, densely packed biofilm clusters and equilibrate at ∼10% of total population, because λ phages only rarely diffuse to areas past a characteristic cell packing threshold and are also neutralized by previously lysogenized cells. Concentrated on the periphery of biofilm clusters, where cell packing is natively low and decreases further due to lytic cycle phage activity (**Supplementary Fig. 7**), lysogens are inherently prone to disperse and become over-represented in the pool of planktonic cells exiting the system. This leads to their overrepresentation in new biofilm populations seeded downstream of the populations from which they originated. The overall nature of temperate phage propagation by host lysis and lysogenization on the biofilm periphery inherently promotes the efficiency of spatial spread of lysogenized cells over the length scale of biofilm dispersal and recolonization.

### Successive phage propagation after lytic induction is blocked by high density cell packing in biofilms

Our work thus far illustrates that while phage λ has limited ability to invade established biofilms of naïve cells, its pattern of lysogenizing hosts along the biofilm periphery leads to overrepresentation in the dispersal pool and frequently a large increase in relative abundance after colonizing new locations. We next asked: in biofilms initially colonized by a mixture of lysogens and naïve cells, does lytic induction of the lysogens allow phages to spread from within and infect hosts that they could not reach when introduced in the liquid phase? To assess this question, we first inoculated biofilms with a mixture of lysogens and naïve host *E. coli* at a ratio of 1:10 (**Figure 2G**). Biofilms were grown for 72 h prior to applying heat to the chambers to induce lysogens to switch to the lytic cycle (See Methods). Note that only a minority of lysogens are induced to lytic phage production. In these experiments, naïve *E. coli* cells carried the constitutive mKate2 reporter, while the inoculated parental lysogens did not; this way, the parental lysogens (mKO2 fluorescent only) could be distinguished from new lysogens (mKate2 and mKO2 fluorescent) produced by phages released from lytic induction events. Following lytic induction, the population dynamics of inoculated lysogens, new lysogens, and remaining naïve cells were tracked daily for 120 h. When we performed this experiment in WT *E. coli* biofilms, we detected negligible production of new lysogens relative to a control case without heat induction **(Figure 3A-C)**. This indicates that the limitation on phage diffusion observed in earlier sections still applies if phages are released from induced lysogens from within established biofilms.

**Figure 3.**
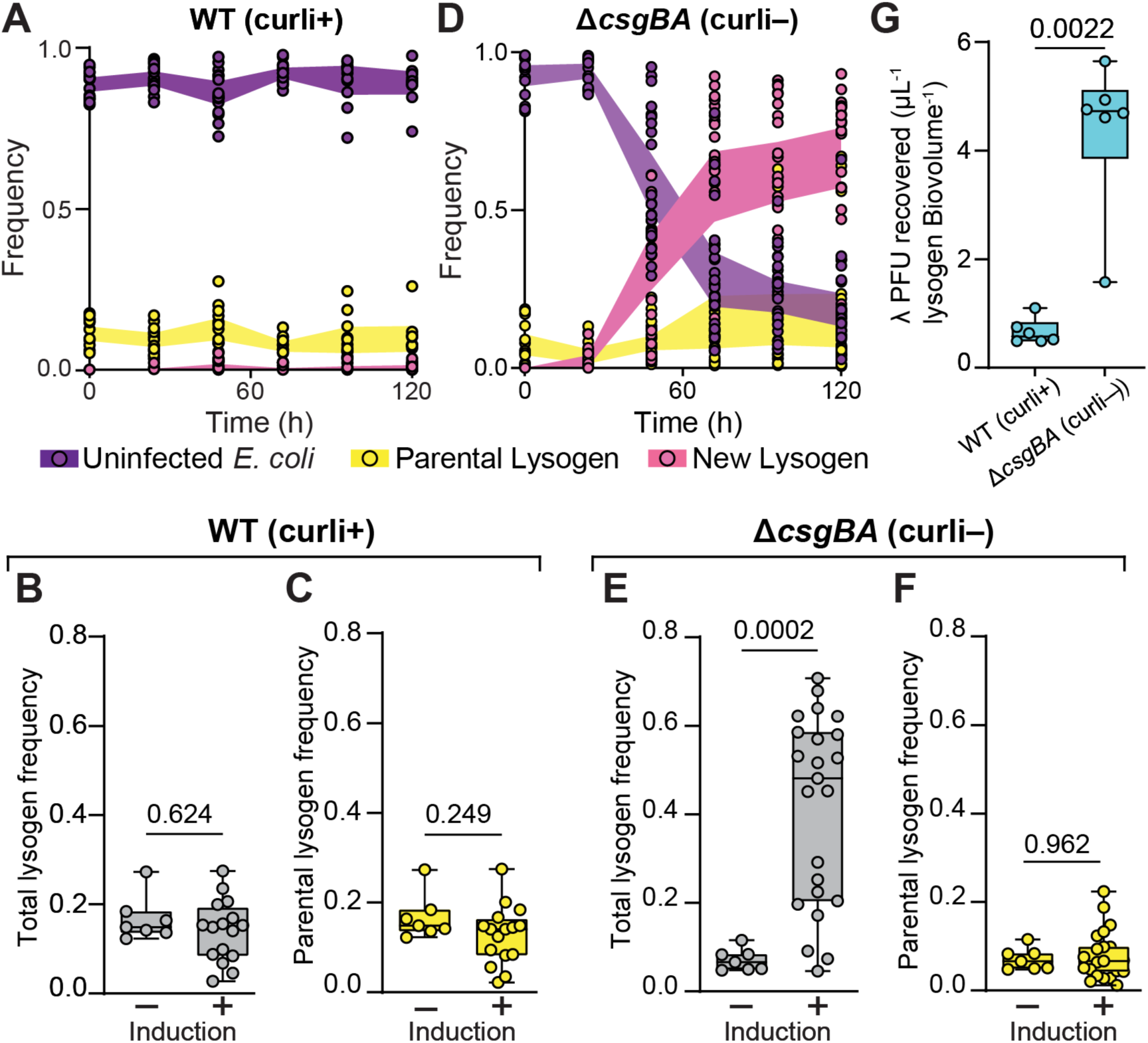
Biofilms of either (A) the WT background were inoculated with a 1:10 mixture of λ lysogens and naïve host *E. coli* and grown for 72 h. Lysogens were then induced to switch to the lytic cycle. The population dynamics were then tracked over 120 h for uninfected *E. coli* hosts, parental lysogens, and newly derived lysogens produced by phages released during the induction treatment at 48 h. (B) Total frequency of lysogens (original inoculated parental lysogens plus newly formed lysogens) and the (C) frequency of only the newly formed lysogens at 120 h in WT biofilms that either had or had not been treated to induce inoculated parental lysogens to switch to the lytic cycle. (D-F) Identical experiments as for (A-C), but in this instance using the Δ*csgBA* curli-null strain background, which does not produce normal cell-cell packing in biofilms and cannot impede phage diffusion. (G) Quantification of phages exiting chambers of WT or Δ*csgBA* biofilms following heat induction to prompt the lytic switch among the lysogens that were co-inoculated with naïve cells.

We speculated that if we again relaxed the cell packing architecture of our experimental biofilms by using the Δ*csgBA* curli null background rather than WT, phages released from induced lysogens may be able to diffuse more freely and infect naïve cells with which the lysogens were inoculated. Supporting this idea, when we inoculated lysogens and naive cells of the Δ*csgBA* background and allowed them to grow before inducing the lysogens’ lytic switch, we observed a steep increase in new lysogen production, with lysogens ultimately reaching the majority of the population by 120 h. (**Figure 3D**). It is interesting to note that the expansion of the lysogen population in these experiments was driven predominantly by the production of new lysogens by released phages, rather than by the growth of the initially inoculated parental lysogens (**Figure 3E,F**). This suggests that time scale of phage transport through Δ*csgBA* biofilms – and the production of new lysogens by infection of exposed naïve cells – is substantially faster than the time scale of further growth by the parental lysogens over the 120 h we tracked the experiments following the lysogen lytic switch induction treatment.

The stark contrast between new lysogen production in WT versus Δ*csgBA* biofilms suggested to us that biofilm architecture coordinated through the expression of curli fibers can block successive rounds of infection and host lysogenization from within a biofilm, even when phages originate from lysogens co-inoculated with naïve cells from the start of the experiment. In the absence of other mechanisms promoting phage mobility through biofilm cell groups, lytic induction of scattered lysogens in the biofilm population contributes little to production of new lysogens in a WT *E. coli* biofilm context. If the mechanism underlying our results is greater phage diffusion through the Δ*csgBA* background, we predict that when lysogens are induced to switch to lytic infection within biofilms of a Δ*csgBA* host, phages should be released into the surrounding liquid more readily than for WT biofilms. We tested this idea by performing the same lysogen induction experiment described above and sampling the liquid effluent of both WT and Δ*csgBA* biofilms for phage titer 3 h after lytic induction of the inoculated lysogens to allow for a full lytic induction cycle to occur at room temperature. We reasoned that if phages have higher mobility through Δ*csgBA* biofilms, they should escape more easily following induction, and we should thus recover higher titers of phage from these biofilms (normalized to lysogen abundance). As expected, we recovered ∼10 fold higher phage titers from Δ*csgBA* biofilms than from WT *E. coli* biofilms **(Figure 3G)** after inducing lysogens to switch to lytic phage production.

### Dissemination of temperate versus obligately lytic phages within and among biofilms

Thus far, we have seen that the temperate phage life history does not necessarily improve within-biofilm phage spread, but because of the spatial pattern of lysogen production in naïve biofilms exposed to phages, phage λ is predisposed to depart into the liquid phase and to reach much higher fractions of the population downstream during recolonization. These observations ultimately prompted us to ask: how do the net benefits and costs of the temperate phage life history for propagation in biofilms compare those of a purely lytic phage? To explore this question, we first performed experiments in which *E. coli* biofilms were grown for 48 h, and then inoculated these biofilms with either phage λ, λΔ*cI*, or T7 phages in separate experimental replicates. T7 is an obligately lytic phage that must kill its host in order to propagate. This strain of T7 contains a transcriptional fusion encoding *sfGFP*, which causes host cells to fluoresce green prior to lysis^31^. We constructed the λΔ*cI* strain by replacing the cI locus encoding Repressor with a truncated nonfunctional copy; this phage retains the mTurquoise2 capsid decoration protein fusion, such that phage particles can still be tracked. cI is required for maintenance of lysogeny in λ lysogens. Without a functional copy of cI for Repressor synthesis, λΔ*cI* can only propagate by lytic infection (**Supplementary Fig. 15**).

We monitored phage propagation in biofilm chambers exposed to phages continuously for 7 h, then removed phages from media flowing into the chambers and returned sterile medium flow to them for 3 h to remove any phages that were not physically associated with the biofilm-dwelling host population. We then connected the effluent lines of these microfluidic devices to new downstream devices and allowed cells and phages to colonize the new chambers for 2 h; finally, new influent tubing was connected to the downstream chambers to introduce sterile media for an additional 96 h. We used infected/lysogenized cell biovolume as our metric for phage spread and propagation in our biofilm chambers, where cells can be determined to be infected if they are undergoing the lytic cell fate or if they are lysogenized. Assessing phage fitness using infected cell counts, including lysogens, has been established as the most robust metric for studying phage population dynamics in recent theoretical work^79–81^.

In the first 7 h of exposure to phage T7, we see the highest accumulation of infected cells just 2 h after phages are inoculated into the system (**Fig. 4A, D**). This is followed by a rapid decrease in infected cells, as the T7 phages kill off most of the cells in the chambers (**Fig. 4A, D**). When we conduct this experiment with λ or λΔ*cI* phages, we observe qualitatively different population dynamics than for T7. Rather than an early peak followed by a subsequent decrease in infected cell biovolume, we see a slow increase in infected cell numbers, contributed to by both lytic and lysogenic infection events by phage λ, and only lytic infection events for λΔ*cI* (**Fig. 4A, C**). If we instead measure the infected cell frequency as a function of the total population, we observe T7 phages reach their highest frequency within 4 h of inoculation in the original chamber, followed by a decrease as cells continue to lyse, exhausting the prey population. Infected cell frequency of λ and λΔ*cI* phages remain low during the 7 h in the original chamber (**Fig. 4B**).

**Figure 4.**
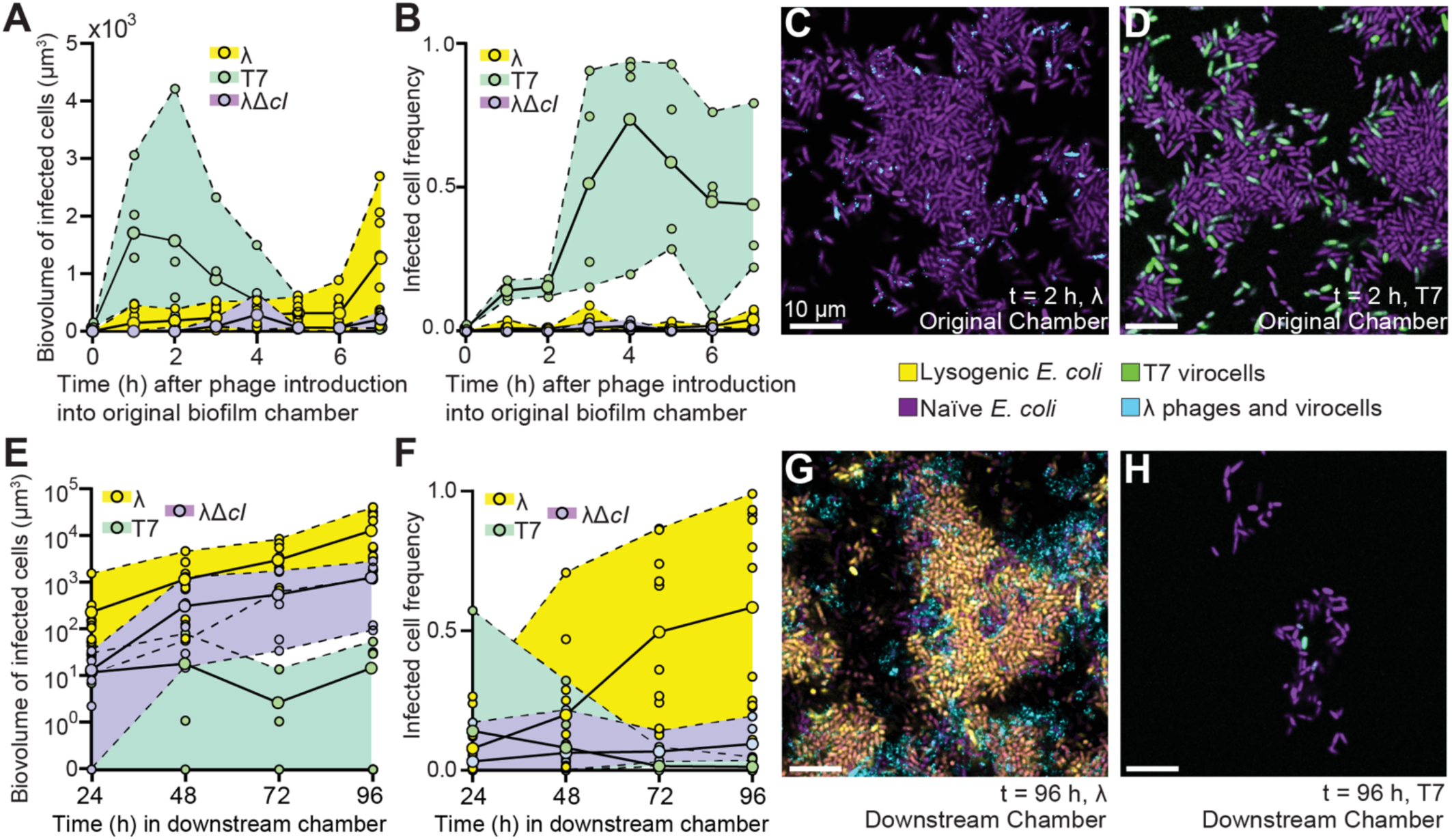
(A) Biovolume of infected cells in original microfluidic devices for 7 h of phage exposure to either T7, λ, or λΔ*cI* phages (n=4-8). (B) Biovolume of infected cells in new (downstream) microfluidic devices over a 96 h time course (n=4-8). (C, D) Representative 2-dimentional optical sections of t = 2 h of exposure to λ or T7 phages, respectively. (E) Infected cell frequency as a proportion of the whole population for T7 or λ in original (upstream) microfluidic device (n=4). (F) Infected cell frequency in new (downstream) microfluidic devices (n=4-8). (G, H) Representative 2-dimensional optical sections of t = 96 h in the new microfluidic devices.

To follow phage and host dynamics over larger spatial scales, we repeated the downstream chamber recolonization experiment in which effluent from original upstream chambers was used to inoculate new, sterile chambers. For phage T7, we occasionally found no cells – infected or uninfected – in downstream chambers (**Fig 4E, H)**. More frequently, we found a small population of *E. coli* cells that had colonized the surface, with a small proportion of these cells actively infected with T7. As a result, the biovolume of infected cells for T7 is lower than for phage λ, and infected cells also make up a small fraction of the total population (**Fig. 4F**). In the downstream chambers exposed to λ phages, we observe an increase in infected cell biovolume, primarily driven by lysogenized cells continuing to divide and populate the new downstream biofilm (**Fig. 4E, G**). The active phage particles in this system also continue to lyse and lysogenize uninfected cells, further contributing to the increase in biovolume of infected cells and high infected cell frequency (**Fig. 4F**). For downstream chambers of λΔ*cI*, we observed a lower infected cell frequency than for λ, but a higher biovolume of infected cells than for T7 (**Fig 4E, F**), which we attribute to T7 having a shorter lag time and higher burst size than phage λ lytic infection events. So, in the original chamber where phages were added continuously for 7 h, we observed T7 accumulating higher numbers of virocells than λ phages. However, in the downstream chambers, we see the opposite effect, where temperate phage λ accumulates 1000-fold higher biovolume of infected cells relative to T7, and λ-infected cells dominate the population over extended time scales. This outcome indicates that virulent and temperate phage strategies are differentially suited to propagation on short time and small spatial scales (for quickly replicating virulent phages), versus propagation over longer time and larger spatial scales (for temperate phages).

## Conclusion

Live, high-resolution imaging of bacteria and phages dwelling in biofilms provides critical insight into the ecology of microbes on the spatial scale at which their behavior directly manifests. Here we tracked the spatial propagation of temperate phage λ in sensitive *E. coli* biofilms, finding consistent patterns in the localization of phages and lysogens along the periphery of biofilm cell groups. Because of diffusion constraints imposed by biofilm cell packing architecture, phages can access only a modest fraction of susceptible *E. coli* in mature biofilms. Furthermore, even once lysogens are present in biofilms alongside susceptible hosts, lytic induction of the lysogens has little impact on promoting new infections, again due to the constraints on phage diffusion imposed by WT biofilm structure. On the other hand, the natural tendency for lysogens to be generated on the periphery of biofilms by phages in the surrounding liquid proved to be important for a different reason. Lysogens’ location on the outer boundaries of otherwise naïve biofilms, where biofilm cell packing is lower and decreased further by lytic phage infections, leads to preferential dispersal of lysogenized cells to new locations downstream of the original site of biofilm growth and phage exposure. Because the temperate phage strategy predisposes itself to dispersal from biofilms of origin, λ phage infection and lysogen production count is far greater over longer time scales and over larger spatial scales on which dispersed cells from one biofilm colonize new downstream populations. This is a novel and unique benefit of the temperate phage life history strategy in comparison with that of obligately lytic phages, and the result emphasizes that examining phage infection dynamics in biofilms can yield new insight into the adaptive benefits of different phage infection strategies that are impossible to infer from traditional shaken liquid cultures.

This study provides a first look into the complex interactions of temperate phages and their host bacteria in a biofilm context. For simplicity, we explored a single species of bacteria and a single strain of phage in any given experiment; further studies can expand on these results by increasing ecological realism in terms of species richness within bacterial biofilm communities, as well as greater diversity among the phages introduced to these communities. While our lysogenized *E. coli* is indistinguishable from parental non-lysogens in terms of biofilm production, many temperate phages change host behavior by introducing novel functions to them via lysogeny, including alterations to metabolism and extracellular product secretion^23,25^. How the traits introduced to their hosts by prophages alter biofilm community dynamics remains a largely unexplored topic and a promising direction for future work. Detailed biophysical modeling of the relative rates of cell growth, cell death due to lysis, phage transport, lysogenic conversion, and cell dispersal will be helpful in defining the conditions under which our observations here are likely to generalize . Lastly, we derived our inferences about the relative benefits and costs of the lytic and temperate phage strategies by growing each separately in different runs of our biofilm experiments. This is an important first step in differentiating how obligately lytic and temperate phages interact with their hosts in the biofilm context; however, in reality, there must be countless scenarios in which temperate and obligate lytic phages directly compete with each other in the same biofilm environments. The question of how temperate phages fare against obligate lytic phages in direct competition for hosts within and among biofilms is another crucial question for future research.

**Table 1.** Reagents, products, and software used in this study.

| Strain | Relevant markers/Genotype | Source |
| --- | --- | --- |
| Bacteria |  |  |
| CNE336 | AR3110, with *csgBAC*-mKate2 transcriptional fusion, *ptac*-mKO-κ and KanR at attB site | <sup>31</sup> |
| CNE859 | AR3110, mTurquoise2/mKO2 $\lambda$, 6x-*His* csgA | This study |
| CNE863 | AR3110, *Lac::mKate2*, 6x-*His* csgA, AmpR-gpD plasmid | This study |
| CNE866 | AR3110, mTurquoise2/mKO2 $\lambda$, 6x-*His* csgA, AmpR-gpD | This study |
| CNE869 | AR3110, *Lac::mKate2*, ΔcsgBA::KanR, AmpR-gpD plasmid | This study |
| CNE881 | AR3110, ΔlamB::KanR, mKate2 | This study |
| CNE905 | AR3110, sfGFP, ΔlamB | This study |
| CNE907 | AR3110, ΔlamB, sfGFP, ΔcsgBA::KanR | This study |
| CNE908 | AR3110, ΔcsgBA::KanR, mKate2, mTurquoise2/mKO2 $\lambda$, AmpR-gpD plasmid | This study |
| CNE909 | AR3110, mTurquoise2/mKO2 $\lambda$, AmpR-gpD plasmid, pSIM5-tetR | This study |
| CNE913 | AR3110, ΔcsgBA::KanR, mTurquoise2/mKO2 $\lambda$, 6x-*His* csgA, AmpR-gpD | This study |
| Phages |  |  |
| CNX11 | T7 with sfGFP under control of phi 10 promoter | 31 |
| λLZ1367 | λD-mTurquoise2 *cl*<sub>857</sub>-mKO2 *bor*::CmR | 66 |
| CNX20 | λD-mTurquoise2, truncated *cl*<sub>1-17::223-239</sub>, *bor*::CmR | This study |
| **Chemicals and reagents** | **Source** | **Product number** |
| Kanamycin | Millipore-Sigma | cat.#60615 |
| MEM Vitamin Solutions | Millipore-Sigma | cat. #M6895 |
| Alexa Fluor™ 633 NHS Ester | ThermoFisher Scientific | cat. # A20005 |
| Anti-6X His Epitope Tag (Rabbit) antibody conjugated to Dylight 405 | Rockland Immunochemicals | cat. # 600-446-382 |
| Poly-dimethylsiloxane (PDMS) | Dow Chemical Company; SYLGARD 184 | cat. # 04019862 |
| #1.5 glass coverslips | Azer Scientific | cat. # 1152260 |
| Inlet tubing | Cole Parmer | cat. # 06417-11 |
| 27Gx1/2 needles | BD Precision | cat. # 30510 |
| 1mL syringes | Brandzig | cat. #CMD2583 |
| Harvard Apparatus Pico Plus Elite syringe pumps | Harvard Apparatus | cat. # 70-4506 |
| **Software and Algorithms** | **Source** | **Version** |
| Zen Black | Zeiss | v14.0.0.0 |
| Zen Blue | Zeiss | v3.4.91.00000 |
| MATLAB | MathWorks | vR2021a |
| Paraview | Kitware | v5.9.1 |
| Prism | GraphPad | v9.4.1 |
| BiofilmQ | (61) | v0.2.2 |

## Acknowledgements

We are grateful to Robert Cramer, Yuncong Geng, Ido Golding, Thu Vu Phuc Nguyen, and Stephen Winans and for feedback on earlier versions of this manuscript, and to members of the Nadell Lab and microbiology community at Dartmouth for feedback on the project. The authors also extend thanks to Thu Vu Phuc Nguyen and Siân Owen for their advice and the pSIM5 plasmid, which were critical for engineering the λΔ*cI* strain used in this study.

## Additional Information

J.B.W. was supported by a NIH T32 training grant (award number AI007519), the John H. Copenhaver Jr. Fellowship, and a GAANN Fellowship. BioMT is supported by NIGMS COBRE award P20-GM113132. CDN is supported by the Human Frontier Science Program (award number RGY0077/2020), the Simons Foundation (award number 826672), NSF IOS grant 2017879, and NIGMS grant 1R35GM151158-01.

## Methods

### Strains

*E. coli* strains used in this project were all AR3110 derivatives. AR3110 was derived from the K-12 strain W3110, which does not produce cellulose because of a polar stop codon mutation in *bcsQ*. This stop codon mutation was corrected in AR3110 to yield a strain that produces cellulose, thus restoring this *E. coli* strain to “wild-type” biofilm formation secreting cellulose and curli protein^69^ . Like other K-12 derivatives, AR3110 is susceptible to both T7 phages and λ phages. Strains of AR3110 were engineered via lambda red recombination or through SacB counterselection allelic exchange. Briefly, primers encoding regions of homology to the flanking regions of the gene of interest were used to amplify the Kan^R^ resistance cassette flanked by FRT sites. For creating fluorescent strains, these resistance cassettes were fused to fluorescent protein constructs via SOE PCR. These PCR products were used to knock out genes of interest and selected for with the resistance cassette. FRT sites were used to flip-out the resistance cassette if desired. SacB allelic exchange was performed by constructing plasmids containing regions of homology to flanking regions and gene insertion, SacB, and a selective resistance marker. Plasmids were transformed into *E. coli* through electroporation and screened for kanamycin resistance, then counter-selected on sucrose plates to remove the antibiotic marker and native allele. Recombinant T7 phages were produced previously using T7select415-1 phage display. Recombinant λ phages were produced by infecting λD*am cI857 bor::KanR* phages on LE392 (permissive host) with the pBR322-λD-mTurquoise2/mNeongreen-E plasmid to allow for recombination, and further selection for fluorescent plaques. λΔ*cI* was created via lambda-red recombination that introduced a truncated nonfunctional copy of *cI* followed by screening for clear plaques^82^. Newly generated strains for this paper will be available upon request.

### Microfluidic flow device fabrication

Microfluidic devices were produced by casting poly-dimethysiloxane (PDMS; Dow Chemical Company, SYLGARD 184, cat. # 04019862) onto premade device molds (a schematic for this can be found in **Supplementary** Fig. 1). The resulting PDMS blocks were cut out of the molds, hole-punched for inlet and outlet channels, and then bonded to #1.5 glass coverslips using plasma cleaning preparation of the PDMS and glass coverslips (Azer Scientific, cat. # 1152260). Between the inlet and outlet port areas, the internal space of the chambers in which biofilms were incubated measured 5,000 μm × 500 μm × 70 μm (LxWxH). Segments of inlet tubing (Cole Parmer PTFE #30, cat. # 06417–11) attached to 27Gx1/2 needles (BD Precision, cat. # 305109) on 1 milliliter syringes (Brandzig, cat. #CMD2583) were plumbed into chamber inlets, and the syringes were pushed by Harvard Apparatus Pico Plus Elite syringe pumps (Harvard Apparatus, cat. # 70–4506). Tubing from chamber outlet channels was fed to effluent waste collection.

### Biofilm culture conditions

Overnight cultures of *E. coli* were inoculated into the microfluidic chambers. After a 45-min incubation period without flow to allow for surface attachment, M9 minimal media with 0.5% maltose continuously flowed into the device at a rate of 0.1 μL/min. For the immunostaining of curli, our AR3110 strain harboring a translational 6xHis tag fused to csgA, which encodes the monomer for curli production, was stained with Anti-6X His Epitope Tag (Rabbit) antibody conjugated to Dylight 405 (Rockland Immunochemicals, cat. # 600-446-382) added to the media at a concentration of 0.1 μg/mL continuously for the entirety of the experiment. Prior work has shown that addition of the 6xHis tag to CsgA does not interrupt its function by any measures tested. All experiments were carried out at room temperature. Note that *E. coli* grows in an open 3-dimensional space in our experiments, and biofilms were imaged (see below) to capture their entire volume. Many representative images in the main text are single 2-dimensional optical sections through the 3-dimensional biofilm image stacks; 2-dimensional images were usually selected for clarity, and to show the biofilm internal activity in addition to the activity on its outer surface area. But note that all biofilms and image data sets refer to the full 3-dimensional system throughout the paper.

### Phage propagation and introduction to biofilms

λ phages were produced by growing lysogenic *E. coli* to an OD600 = 0.2 in M9 minimal media with 0.5% maltose, then heat shocked at 42°C for 20 min, and then incubated at 37°C until visible lysis occurred. T7 phages were produced by growing sensitive E. coli to OD600 = 0.4 in M9 minimal media with 0.5% maltose, before adding an aliquot of T7 phage and incubating until the bacterial cultures were cleared. Phages were quantified using standard plaquing techniques and back-diluted to 10^4^/μL in M9 with 0.5% maltoseTo visualize T7 phage infection, we used a previously constructed T7 strain that induces sfGFP production by the host prior to lysis. For experiments with short phage exposure, phages were continuously introduced for 24 h. For experiments with extended phage exposure, phages were continuously added for 120 h. For the nascent biofilm phage exposures investigating spatial patterning in the absence of established biofilms, phages were introduced into the chamber immediately following the initial attachment step.

### Biofilm dispersal and detection of *de novo* λ-resistance mutants from biofilm culture

*E. coli* biofilms grown for 48 h and then treated with λΔ*cI* phages for 24 h were dispersed from the biofilm by removing the tubing from the microfluidic device and vigorously pipetting 100 μL of M9 media and air bubbles back and forth between the inlet and outlet ports. This was done to ensure maximal removal of all cells in the chamber in order to capture accurate measurements of total *E. coli* and any de novo λ-resistant *E. coli*. To determine cell viability and phage sensitivity, this 100 μL volume containing dispersed biofilm cells was serially diluted and plated on LB plates for total *E. coli* counts and, in parallel, on LB plates saturated with λΔ*cI* phages to determine *de novo* λ-resistant counts.

### Phage adsorption assay

Bacterial cultures of uninfected *E. coli*, lysogenic *E. coli*, and *E. coli ΔlamB* were grown and back-diluted to an OD600 = 0.2 in λ medium. λ phages were added to a final concentration of 5×10^3^ phages per μL. Cultures were incubated at 30°C on an orbital shaking platform, and 200 μL aliquots were taken every 25 min, passed through a 0.2-μm filter, and stored on ice until the end of the experiment. The filtration step served to exclude any bacterial cells, and any phages that were attached to them, allowing us to measure free phages remaining in the liquid medium. Flowthrough samples were then serially diluted and plated for PFUs.

### Phage adsorption and *E. coli* population dynamics assay

Sensitive *E. coli* was grown and back-diluted to an OD600 = 0.2 in λ medium. λ phages were added to a final concentration of 5×10^3^ phages per μL. Cultures were again incubated at 30°C on an orbital shaking platform, and 200 μL aliquots were taken every 25 min, passed through a 0.2-μm filter, and stored on ice until the end of the experiment. Another 20μL aliquot was taken and directly added to a dilution series in a 96-well plate. To avoid *E. coli* amplification after sampling, these samples were immediately serially diluted and plated for CFUs. After all of the samples were collected, flowthrough phage samples were serially diluted for PFUs. To determine phage adsorption onto lysogenic cells within biofilms in the absence of curli production, biofilms of uninfected cells and lysogenic cells in a Δ*csgBA* genetic background were grown at a 1:1 ratio for 48 h prior to phage addition for 2 h. Biofilms were then imaged for phage localization around the two strains. This experiment was also performed with uninfected cells and Δ*lamB* cells, which do not have the λ phage receptor.

### Biofilm effluent and recolonization assay

Biofilms were incubated for 48 h without phage exposure, treated with phage for 24 h, and then imaged. We then shortened the effluent tubing to allow for effective collection, and increased flow to 1 μL/min to collect 10uL of media. A portion of this sample was imaged under an agar pad to determine lysogen abundance. Another portion of this sample was used to inoculate new microfluidic devices, incubated without the addition of exogenous phages. These new microfluidic devices were tracked through time.

### Lysogen induction assay

Naive, phage-sensitive *E. coli* and lysogenic *E. coli* were inoculated into biofilm chambers at a ratio of 10:1 for 72 h. Our λ phage carries the heat-sensitive *cI*857 allele; to induce lysogens to convert to the lytic cycle, microfluidic devices were placed into a 42C° incubator for 40min. This was sufficient to induce some, but not all, of the lysogenized cells to switch to lytic phage production. Biofilms were then tracked for 120 h to measure the population dynamics of sensitive cells, parental lysogens, and new lysogens. New lysogen lineages – i.e., those that were created within the biofilm by phages released by the initially inoculated lysogens – can be differentiated from parental lysogen lineages, as new lysogens produce two different fluorescent proteins (mKate2 and mKO2) while parental lysogens only produce one fluorescent protein (mKO2). This, however, does not allow for differentiation between novel lysogenic infections and asexual bacterial replication or newly lysogenized hosts. In order to identify how biofilm structure is important for phage propagation, this experiment was carried out in a WT *E. coli* biofilm background, as well as a *ΔcsgBA* genetic background that cannot produce curli matrix proteins.

### Phage mobility assay

In order to determine phage mobility on a global scale, Biofilms of lysogenic *E. coli* and Δ*lamB E. coli* in a WT background and *ΔcsgBA* background were grown at a ratio of 10:1 for 72h, quantified for lysogen biovolume, and lysogens were induced at 42C° for 40min. Effluent was collected from the microfluidic devices and PFUs were quantified and normalized to lysogen abundance in the biofilm chambers.

### Microscopy and image analysis

All imaging was performed using a Zeiss 980 line-scanning confocal microscope, using a 40x/1.2 N.A. water objective or a 10x/.4 N.A. water objective. The 6xHis Tag Antibody Dylight 405 that was used to stain 6xHis-tagged curli polymers was excited with a 405 laser line. The mTurquoise2 protein that λ and λΔ*cI* phage capsid produces was excited with the 458 laser line. The sfGFP protein produced by the T7 infection reporter construct was excited with a 488 laser line. The mKO2 protein that lysogenic *E. coli* expresses constitutively was excited with a 543 laser line. The mKate2 protein that WT *E. coli* and Δ*csgBA E. coli* express constitutively was excited with a 594 laser line (in separate experiments). If one image was not sufficient to capture the variation seen within a given microfluidic device, multiple independent locations were chosen within each biofilm chamber and averaged to give 1 biological replicate measurement for a given chamber in the case of whole-biofilm measurements. Prior to export, images were processed by constrained iterative deconvolution in ZEN blue. Raw image data was then exported from Zen to the Biofilm Q image processing framework. Constitutive reporters for marking different strains, phages, and lysogens, were binarized using Otsu or Robust Background thresholding with a manual sensitivity parameter. For all analyses, a 3-dimensional grid was used to partition the segmented biovolumes into pseudo-cell cubes that were 0.72 μm on a side. Cell packing measurements merged the biovolume of all bacteria within a sample and calculated the biovolume fraction within 6 μm of each segmented bacterial volume within each grid cube. To measure *csgBAC* transcription for Supplemental Fig. 10, we quantified the *csgBAC* fluorescent reporter fluorescence inside each segmented volume of *E. coli*. To measure curli protein immunostaining, we measured curli immunofluorescence in a 0.5 μm extending away from the outer surface of every segmented volume of *E. coli*.

### Replication, quantification, and statistics

Replication is reported for each experiment individually in the legends of all of the figures. The reported sample size for each figure panel refers to biological replicates. One biological replicate was defined as the averaged outcome for measurements across a single microfluidic flow chamber inoculated from independent overnight culture preparations. Biological replicates for the core experiments in the study were performed across 3 weeks with independent microfluidic chambers. Technical replicates were separate z-stacks captured at randomized locations throughout a given flow chamber; measurements from these technical replicates were averaged to calculate the value for the biological replicate corresponding to that flow chamber. Mann–Whitney U tests with the Bonferroni correction were used for pairwise comparisons. We chose nonparametric comparison tests because they are relatively conservative and because the assumptions required for parametric tests could not consistently be assessed for our data. Boxplots denote the median, interquartile range, and range limit values; individual data points are shown where possible as well.

## Supplemental Information

**SI Figure 1.**
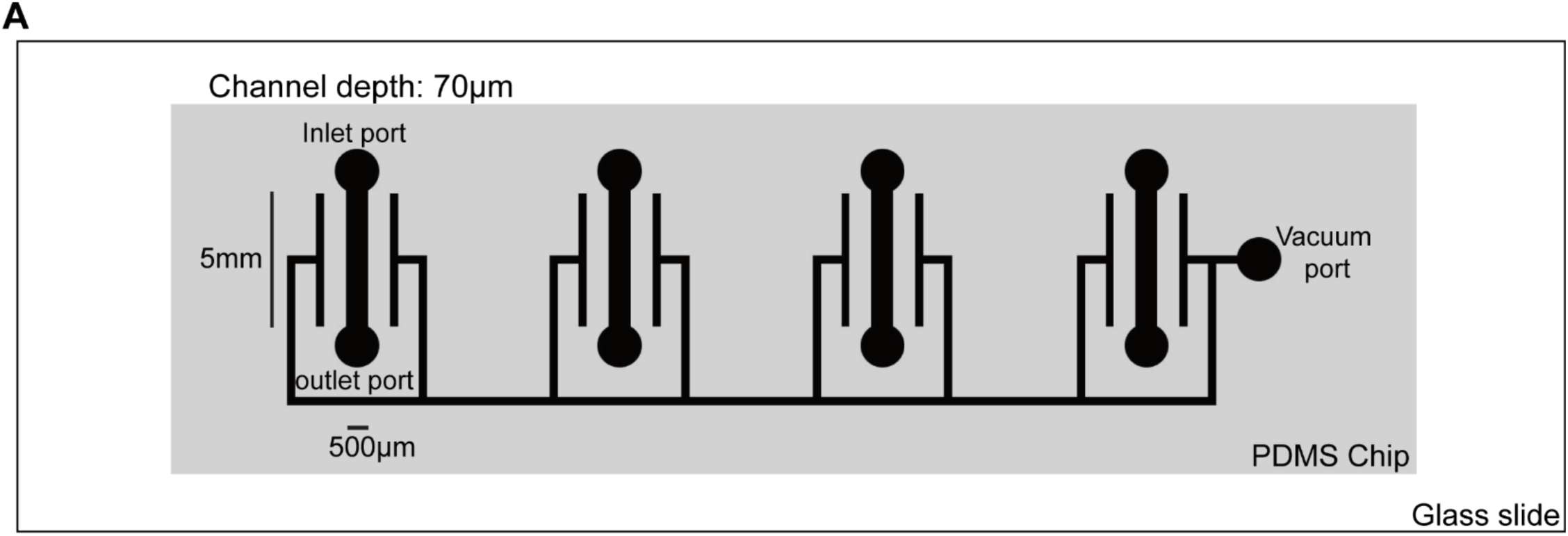
Diagram of our microfluidic devices. (A) This example contains 4 parallel chambers, each with an inlet and an outlet port for connection to fluid inlet/outlet tubing. Technical replicate image stacks were taken from within the straight rectangular section between the rounded inlet and outlet ports of a chamber. The thinner, continuous channel surrounding the 4 separated chambers was connected to a wall vacuum line to apply negative pressure; this method discourages the introduction of air bubbles into the liquid-filled portions of the flow chambers.

**SI Figure 2.**
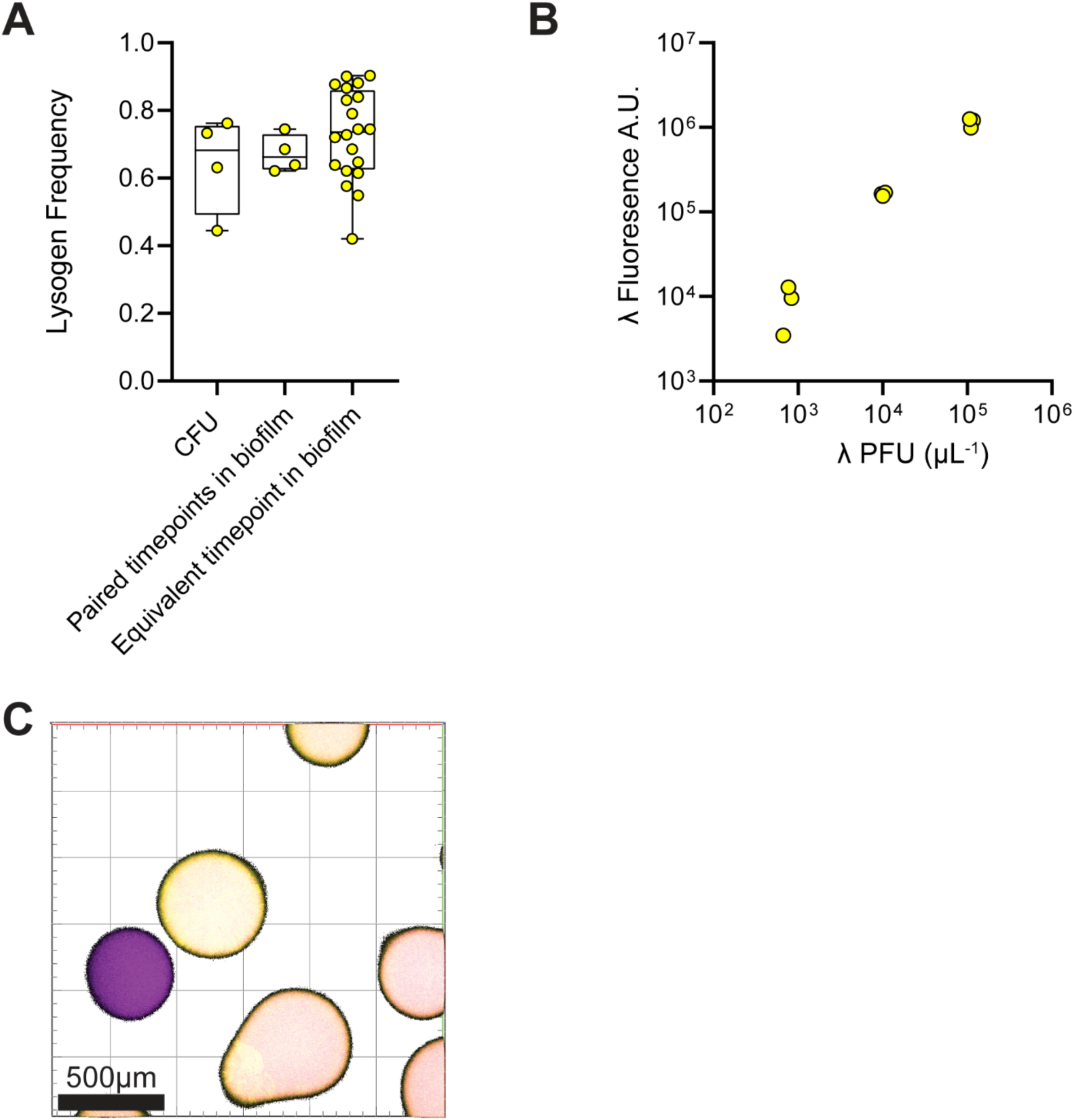
(A) Lysogen frequency measured by CFU selective plating after mass dispersal of biofilms from the microfluidic devices, compared to the lysogen frequency observed in the microfluidic devices measured by image quantification with BiofilmQ (n = 4–20). (B) Linear relationship between λ phage fluorescence and λ PFU (n=9) (C) Lysogenic *E. coli* colonies on LB agar plates can be distinguished from naïve *E. coli* colonies by fluorescence.

**SI Figure 3.**
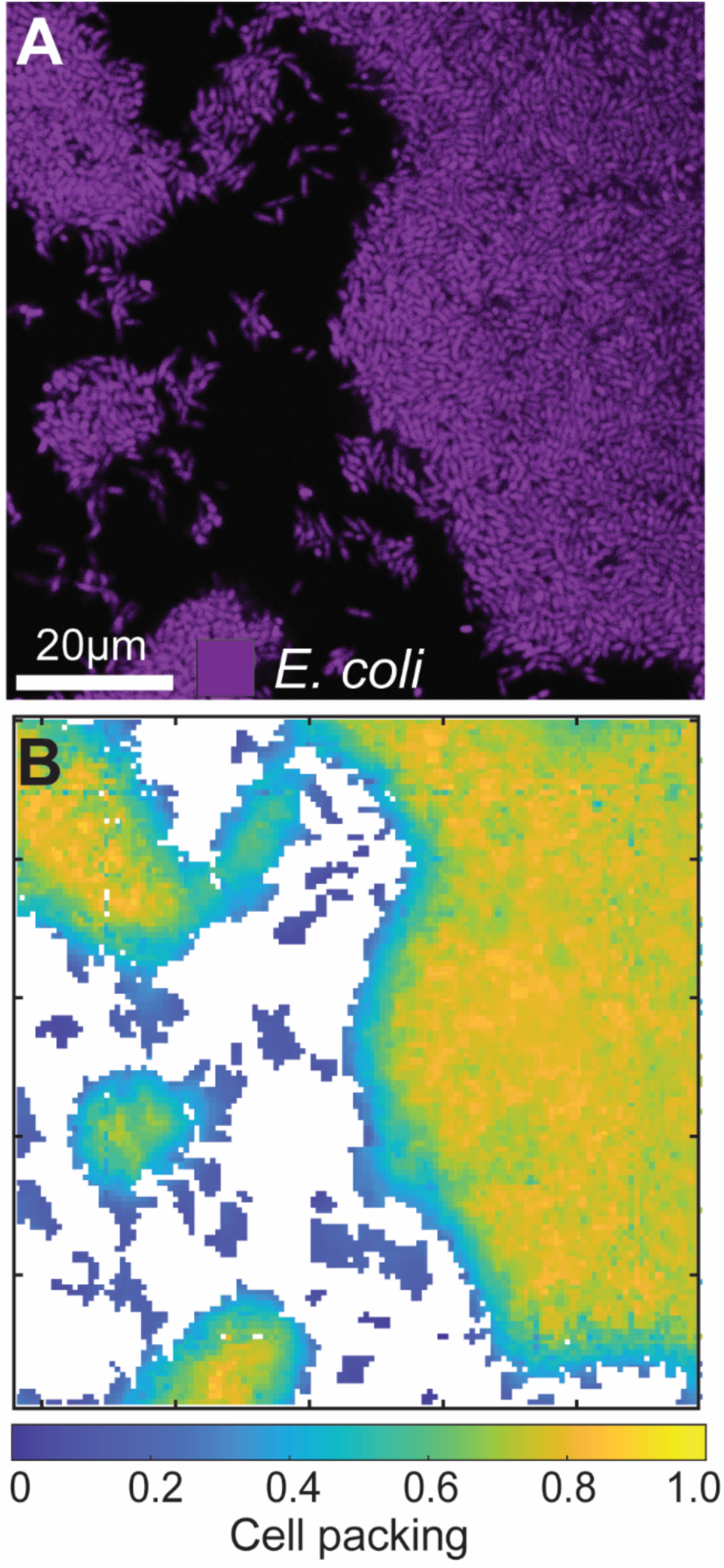
(A) Representative image of an *E. coli* biofilm prior to the addition of λ phages. (B) Heatmap of localized cell packing of (A), showing the variation often seen in *E. coli* biofilm cell packing, dependent on the elapsed time of growth before exposure to phages. The heatmap is a 2-D projection of the analysis for the full 3-D z-stack of images captured.

**SI Figure 4.**
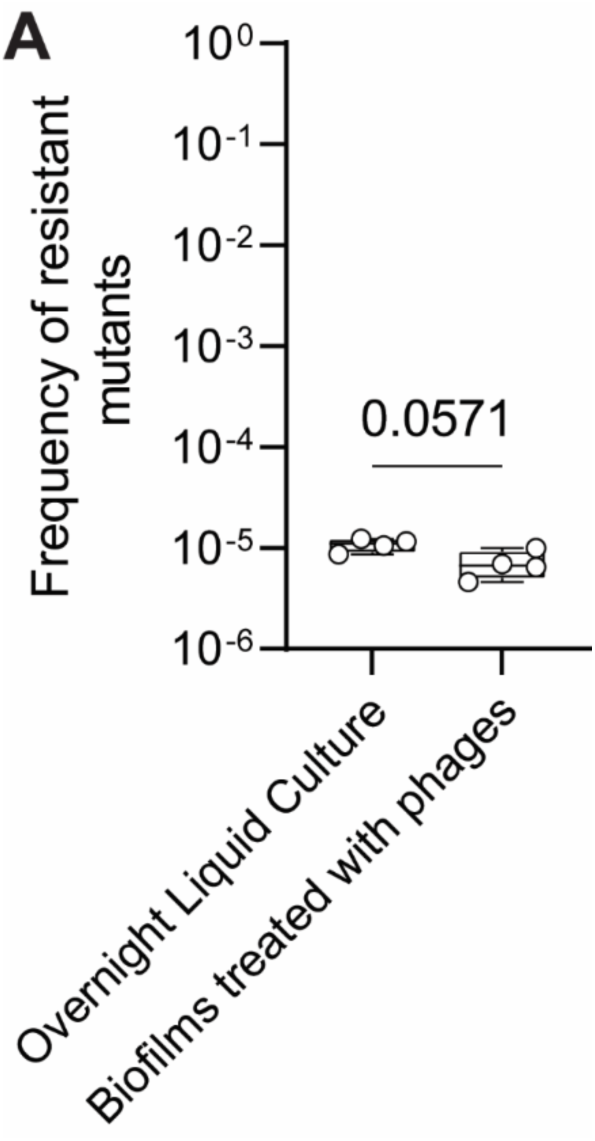
*E. coli* was grown for 16 h in LB media at 37C° overnight. We then plated for CFUs in the absence of phage, and simultaneously incubated *E. coli* with λΔ*cI* phages for 30minutes prior to plating onto soft agar. We then grew *E. coli* biofilms for 48 h, treated them with λΔ*cI* phages for 24 h, then flushed the chambers as described in (Methods). We then incubated *E. coli* with λΔ*cI* phages and plated onto soft agar for resistant colony formation. We found no difference in the frequency of resistant mutants.

**SI Figure 5.**
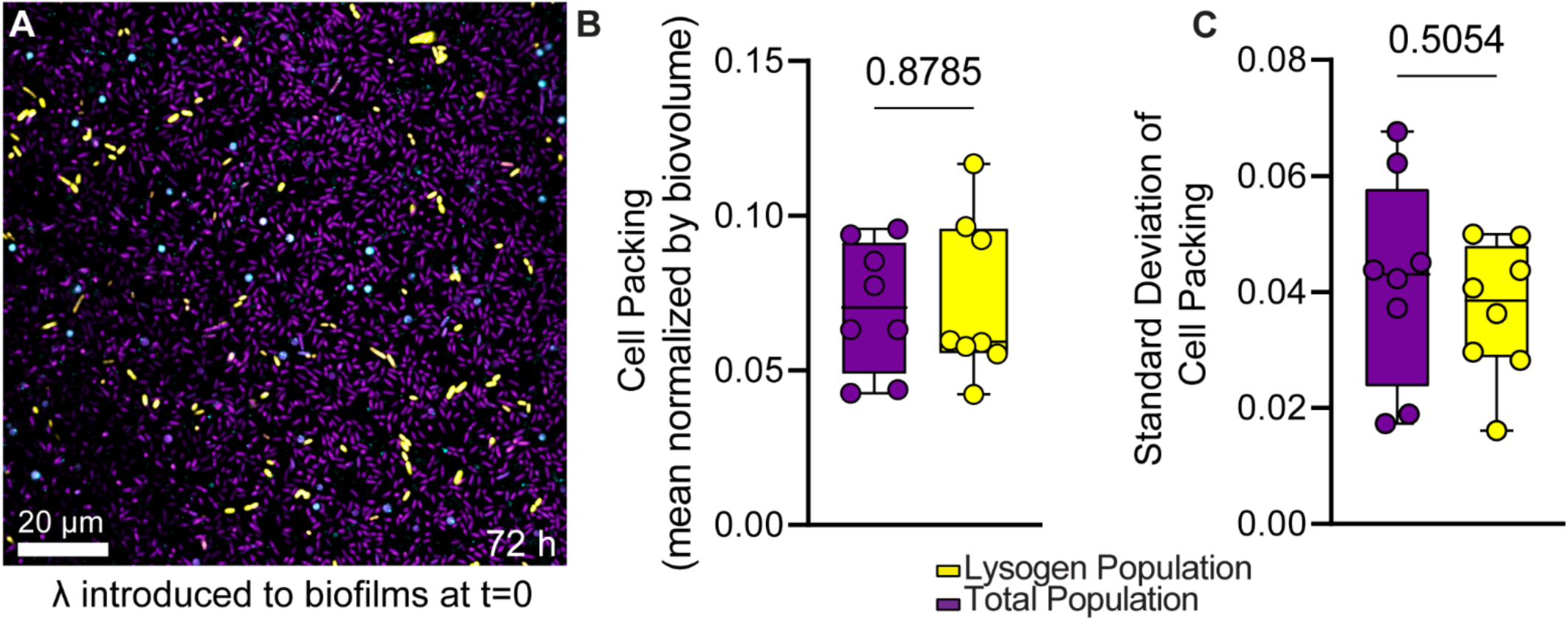
(A) λ phages introduced to *E. coli* biofilms at t=0 (i.e. at the same time as bacterial inoculation) do not exhibit the characteristic pattern of phage spread and lysogenization as observed in established WT *E. coli* biofilms. (B) Cell packing of the total population compared to the lysogenic population, which are not significantly different (n=8). (C) The standard deviation of cell packing for the total population versus lysogenic population, also not significantly different (n=8).

**SI Figure 6.**
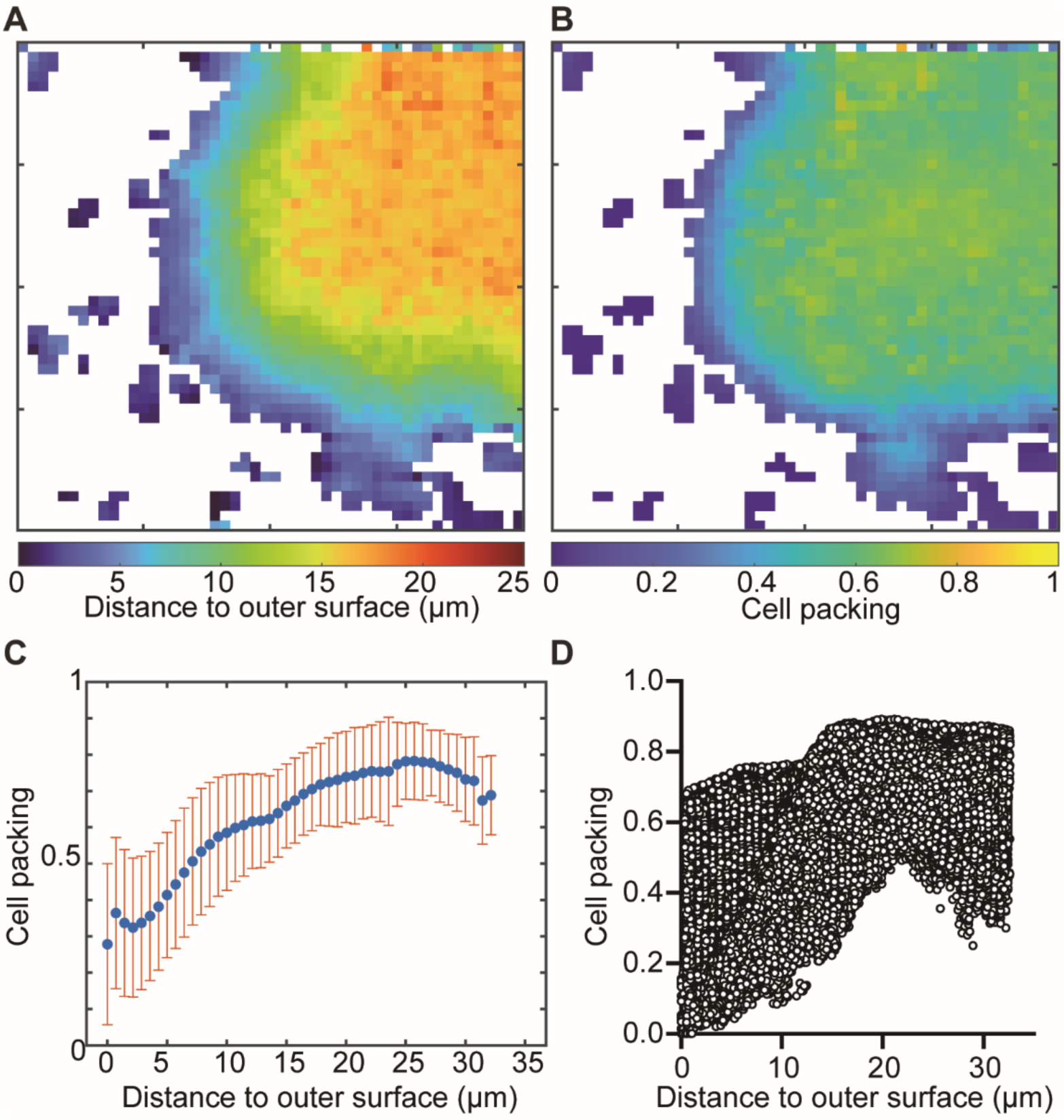
(A) Representative heatmap of distance to outer surface (µm). (B) Representative heatmap of cell packing for the same image data as (A). (C) 1.5D Histogram of cell packing as a function of distance to outer surface (n=9). (D) Plot of cell packing as a function of distance to outer surface for each segmented volume of *E. coli* in replicate image data sets. Note that segmented bacterial volumes are partitioned into a 3-dimensional grid to delineate cell-sized objects (which in this case, are the volumes within each cubic node in the 3-dimensional grid) and to create a reference frame for spatial analysis (n=9, total segmented volumes of *eE. Coli* represented from the 9 image sets =234,695).

**SI Figure 7.**
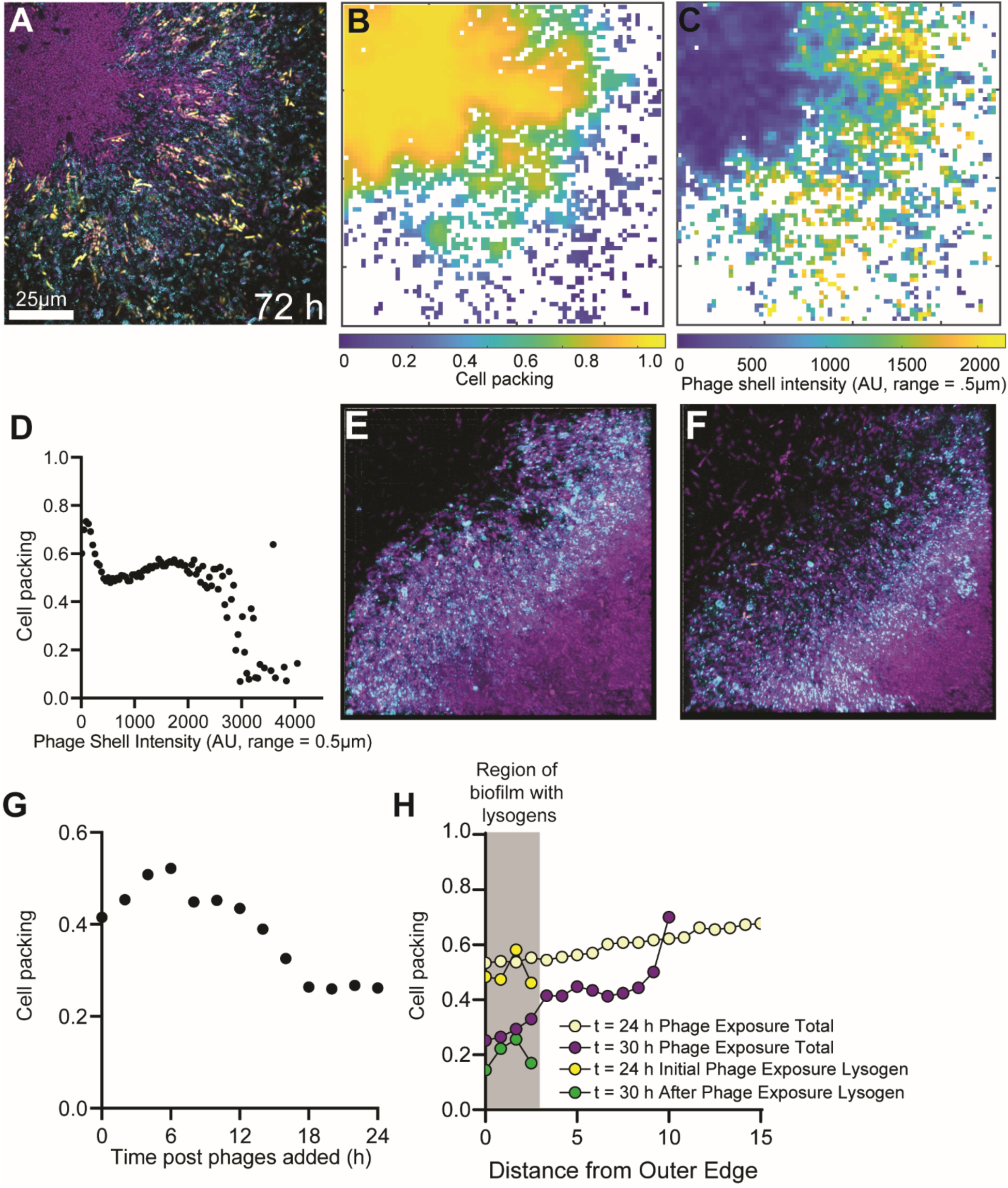
(A) Representative image of an *E. coli* biofilm exposed to phages for 72 h. Naïve *E. coli* hosts are shown in purple, phages in cyan, and lysogens in yellow. (B) Corresponding heatmap of cell packing for the biofilms shown in panel (A). (C) Corresponding phage shell fluorescence intensity around each segmented *E. coli* volume within a 0.5μm radius. (D) Cell packing as a function of phage shell intensity, showing a negative relationship. (E) First time point of a high resolution timelapse t=24 h (93µm x 93µm x 15µm), (F) Last time point of a high resolution time lapse t=30 h (93µm x 93µm x 15µm).(G) Data from a high resolution timelapse showing changes in cell packing 24 h after phages have been introduced to the biofilm. Cell packing decreases over time due to phage activity on the periphery of biofilm clusters. (H) Quantified data from the timelapse in (E) and (F), showing cell packing as a function of distance from the outer periphery of the biofilm before and after phage exposure. We observed a decrease in cell packing after phage exposure on the periphery of the biofilm where lysogens are formed.

**SI Figure 8.**
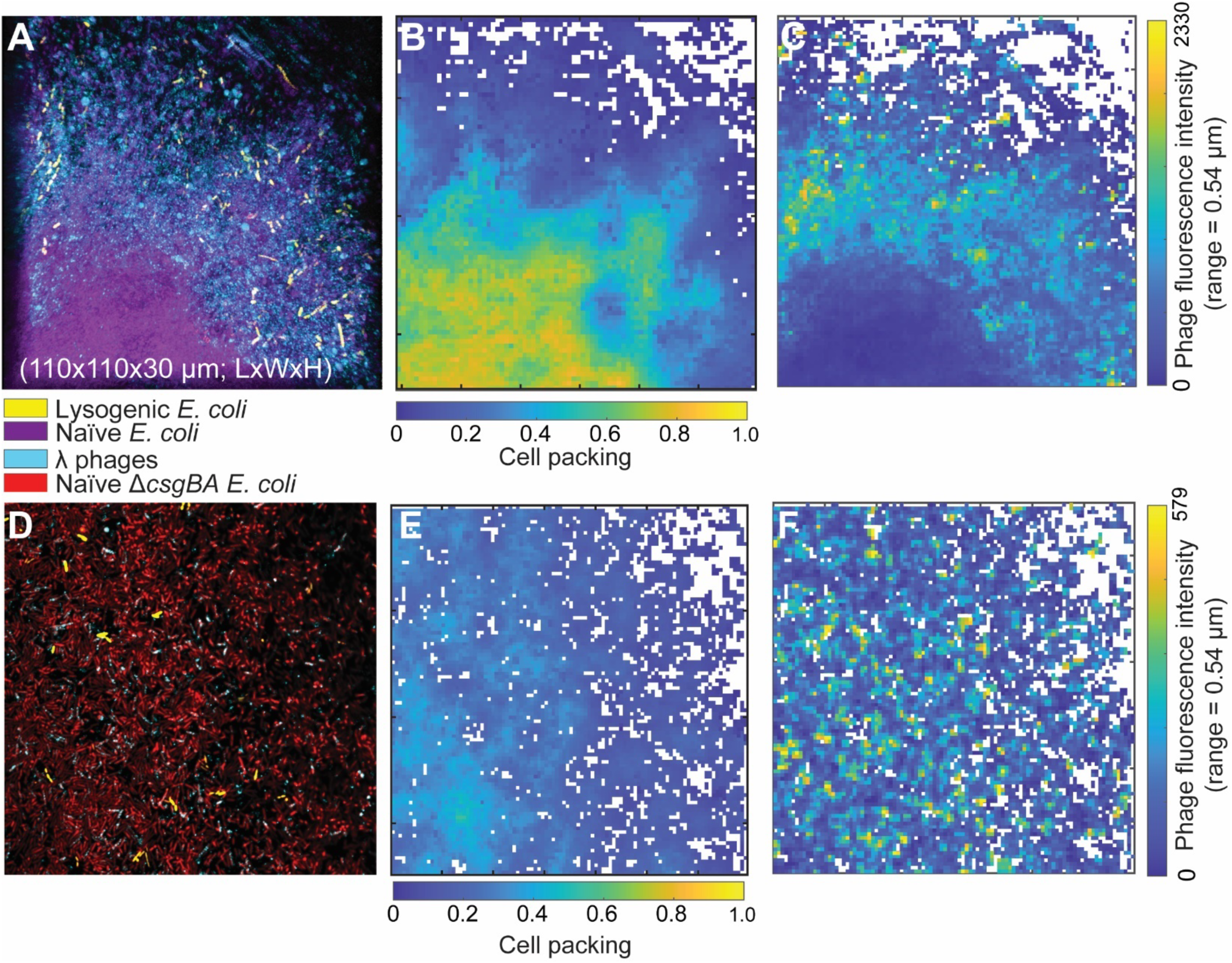
(A) WT *E. coli* biofilm invaded with λ phages for 24 h. (B) Heatmap of cell packing of the established biofilm. By visual inspection, the lysogens aggregate in a region of medium-low cell packing. (C) Heatmap of phage fluorescence intensity surrounding *E. coli.* (D) Δ*csgBA E. coli* biofilm invaded with λ phages for 24 h. (E) Heatmap of cell packing of the established biofilm, with little spatial patterning of lysogenic cell aggregation. (F) Heatmap of phage fluorescence intensity surrounding *E. coli*.

**SI Figure 9.**
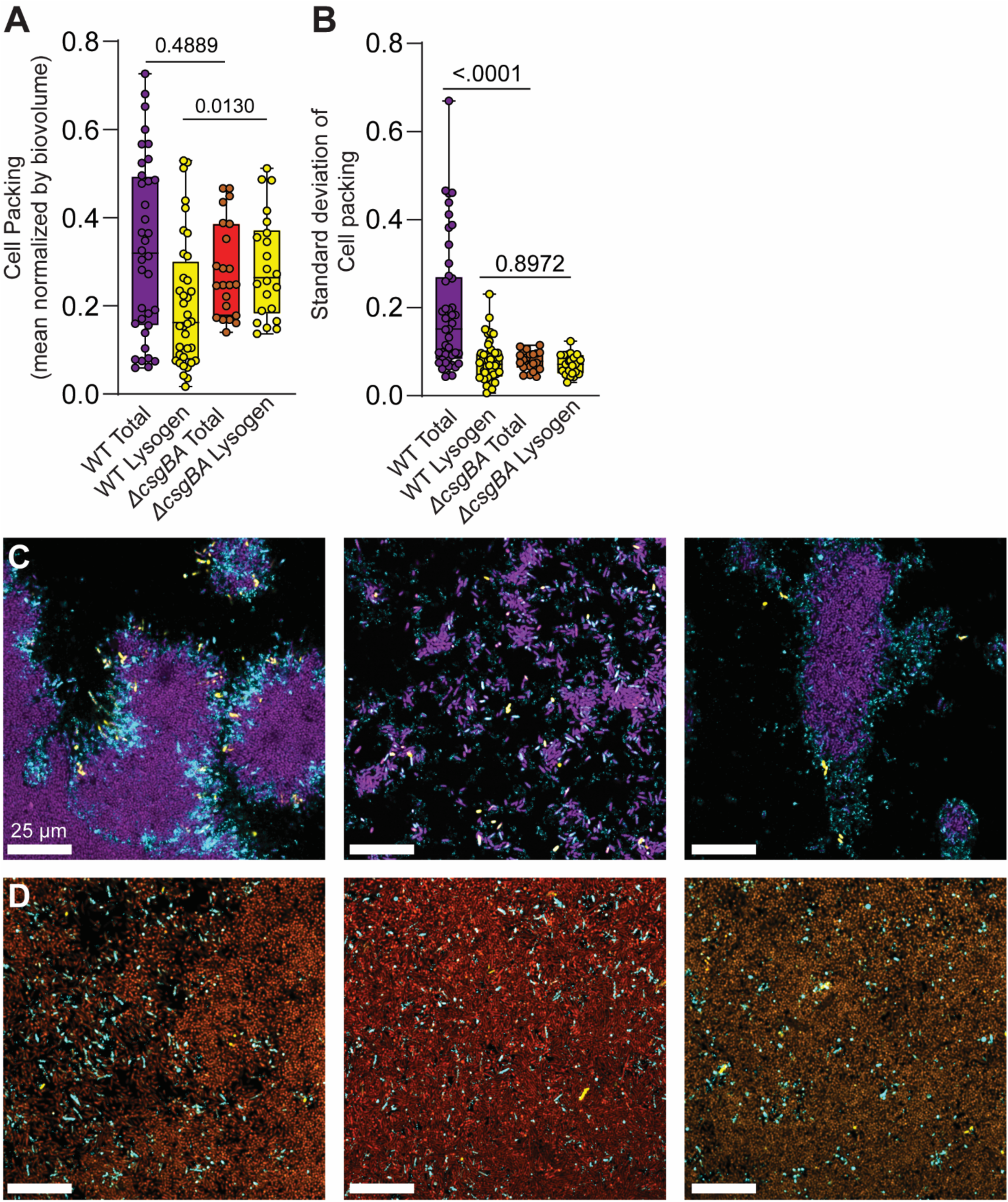
(A, B) Quantative analysis of the mean cell packing and standard deviation of cell packing (as in Figure 1 E and F) showing no significant difference between the mean cell packing of WT and Δ*csgBA* biofilms, but a significant difference in mean cell packing between the lysogen population formed in WT biofilms, relative to those formed in Δ*csgBA* biofilms. We observe a significant difference between the standard deviation in cell packing for the total WT and Δ*csgBA* biofilm populations, but not a significant difference between the standard deviation in cell packing around their respective new lysogen populations. (C) Three independent representative images of WT *E. coli* biofilms (purple) exposed to phages (cyan) for 24 h with novel lysogen infections arising (yellow). Representative images of Δ*csgBA* biofilms (red) undergoing the same phage exposure experiment, again with phages in cyan and lysogens in yellow.

**SI Figure 10.**
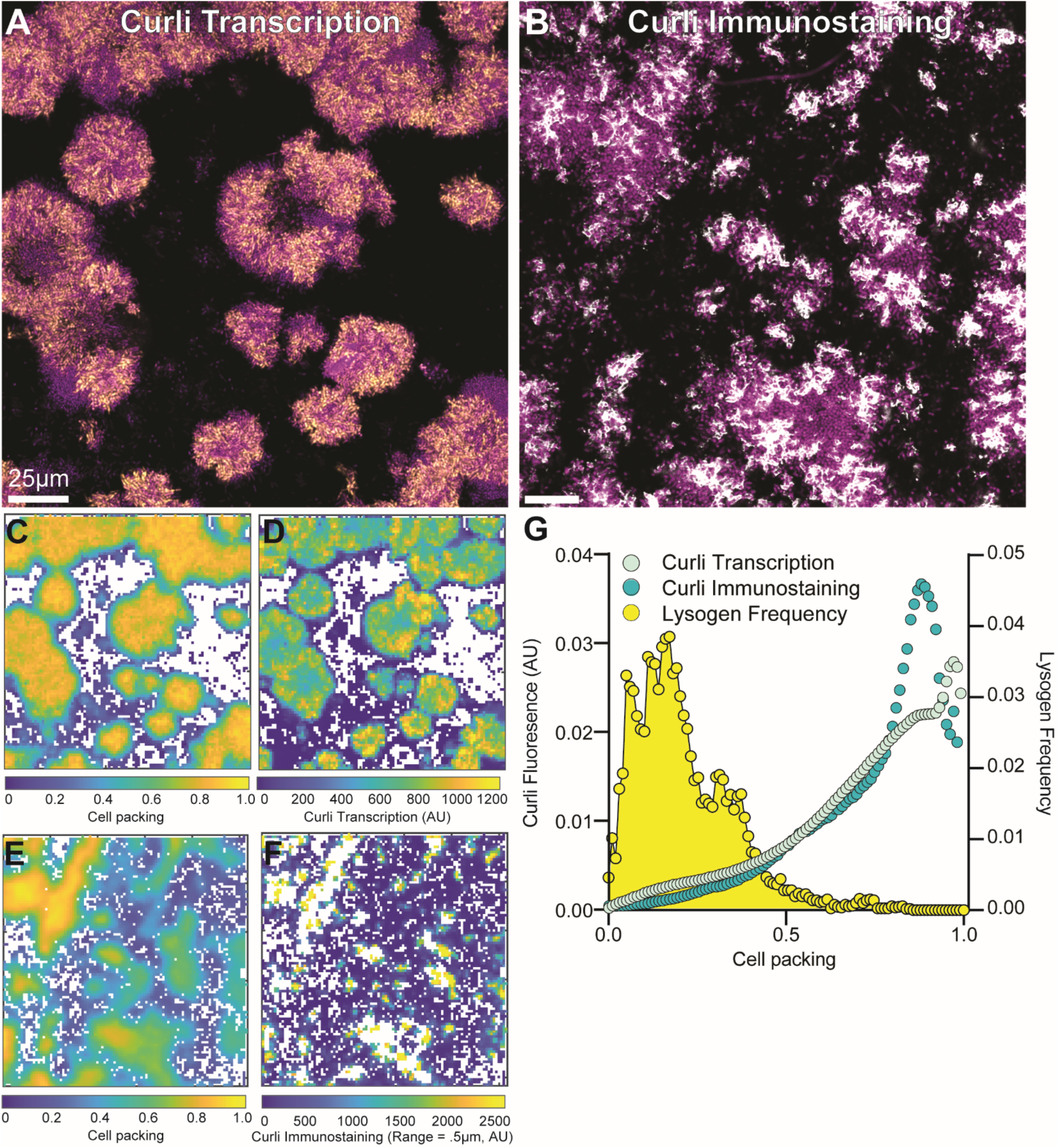
(A) Representative 2-dimensional cross section of *E. coli* (purple) containing the transcriptional reporter for the *csgBAC* operon (fluorescence from the reporter is shown in yellow). (B) Representative 2-dimensional cross section of of *E. coli* (purple) with fluorescent immunostaining of CsgA-6xHis produced from the native genomic locus (immunofluorescence shown in white). (C) Cell packing of biofilms shown in panel (A). (D) Curli transcription heatmap of panel (A). (E) Cell packing of biofilms shown in panel (B). (F) Shell fluorescence intensity with a range of .5μm around *E. coli* cells, staining for curli-associated immunofluorescence. (G) Curli transcription and immunostaining fluorescence intensity as a function of Cell packing from independent replicates (n=12, n=8, respectively). Lysogen frequency as a function of biofilm cell packing (yellow distribution) denotes the same data as used in Figure 1 of the main text.

**SI Figure 11.**
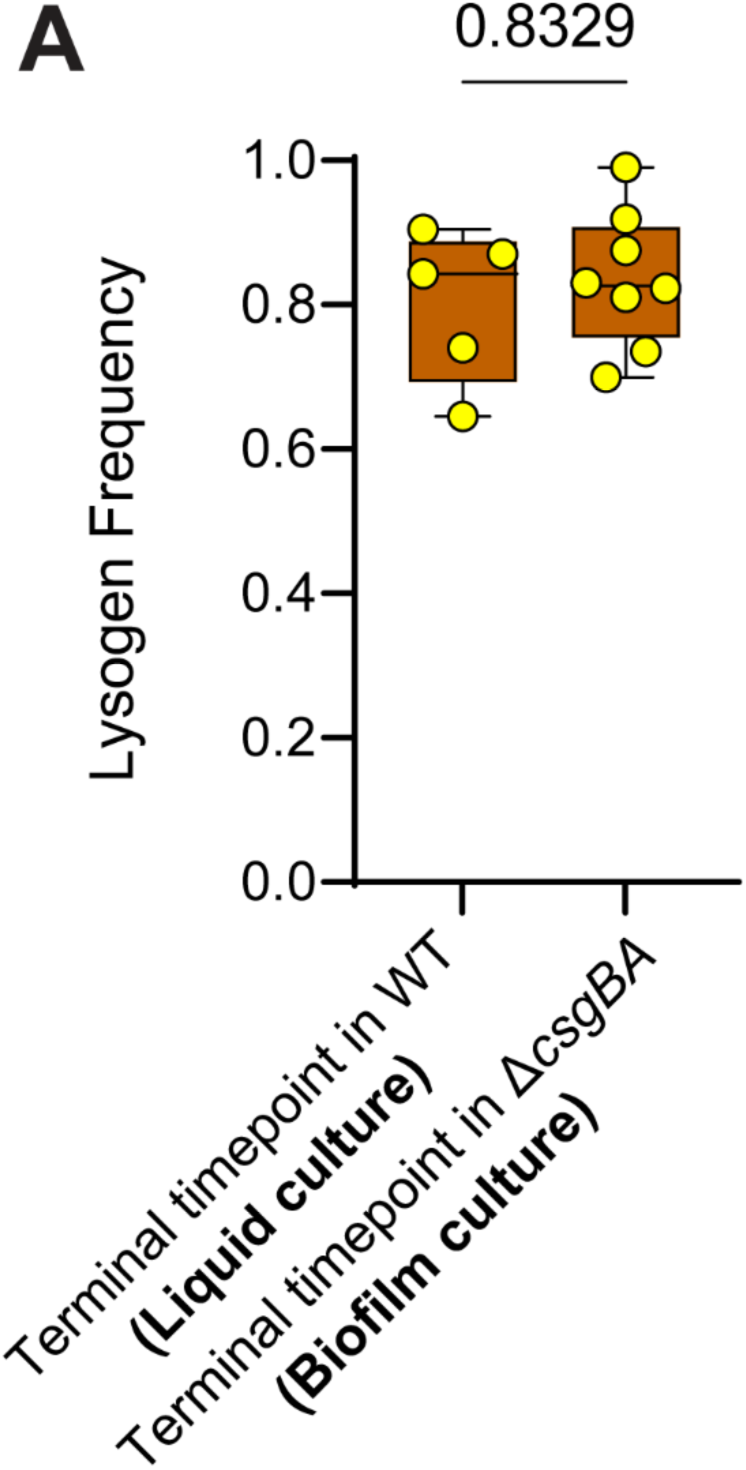
(A) Lysogen frequency within our well-mixed liquid culture conditions was not significantly different from our lysogen frequency in microfluidic device culture of Δ*csgBA* biofilms (n = 5–9). P-values are shown for Mann-Whitney U-tests for pairwise comparison.

**SI Figure 12.**
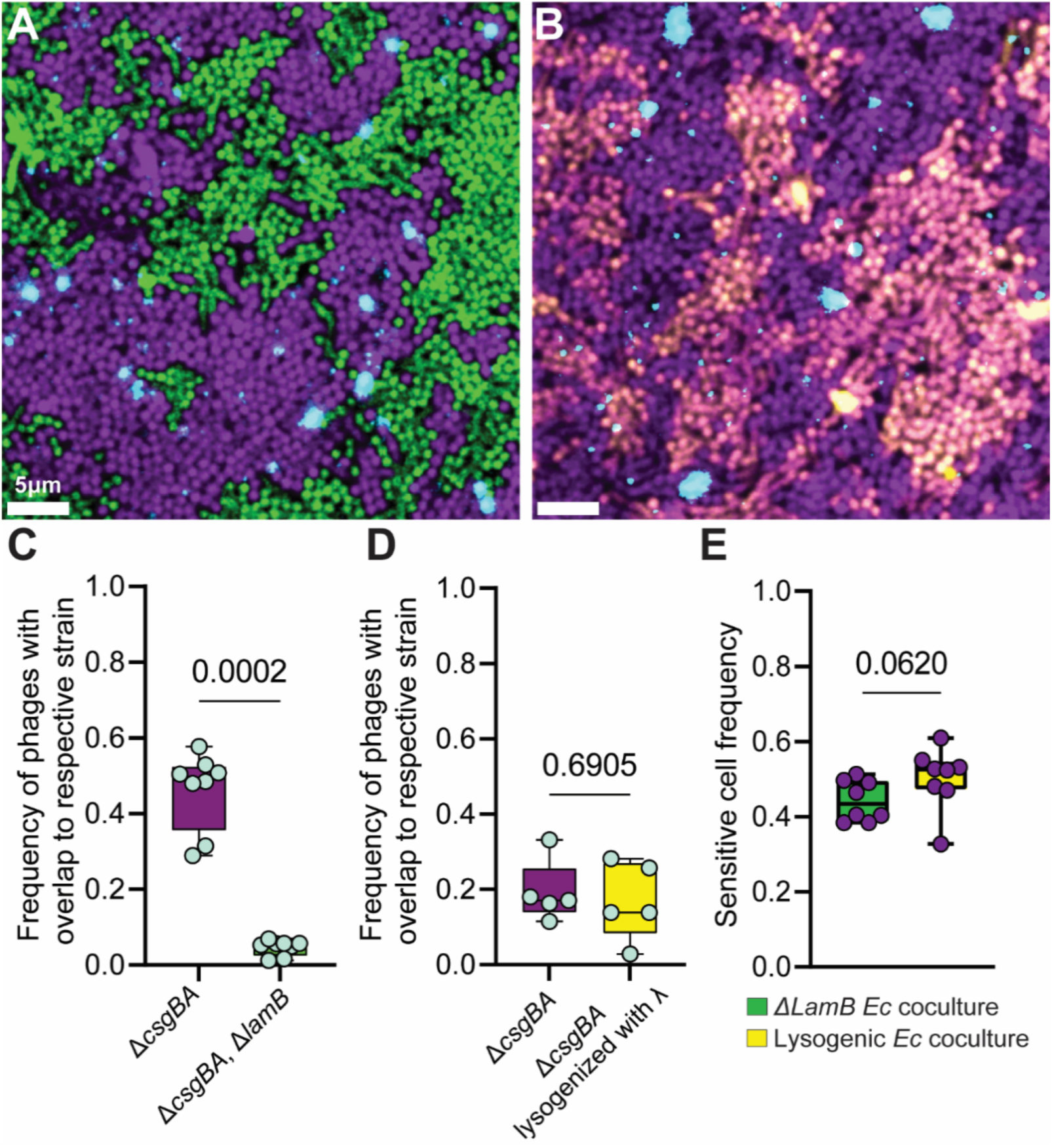
These experiments were designed to determine if phage adsorption/neutralization by lysogens could lower the exposure of nearby susceptible cells to phages. To study this question, we aimed to remove the effects of impeded phage diffusion due to biofilm architecture, and so the experiments are performed in a Δ*csgBA* curli-deficient background, which permits phage diffusion freely relative to the WT genetic background. (A) Coculture biofilms of of Δ*csgBA* (purple) and Δ*csgBA,* Δ*lamB* (green), inoculated with λ phages (Turquoise) for 24 h. (B) Coculture of Δ*csgBA* (purple) and Δ*csgBA* lysogens (yellow) with phage λ (cyan), inoculated with λ phages for 24 h. (C) Frequency of phages with positive Manders’ overlap for Δ*csgBA* and Δ*csgBA* and Δ*lamB* (n=8) (D) Frequency of phages with positive Manders’ overlap for Δ*csgBA* and Δ*csgBA* lysogenized with λ (n=5). (E) Sensitive cell frequency in the two separate coculture experimental setups (n=8). P-values are shown for Mann-Whitney U-tests for pairwise comparison.

**SI Figure 13.**
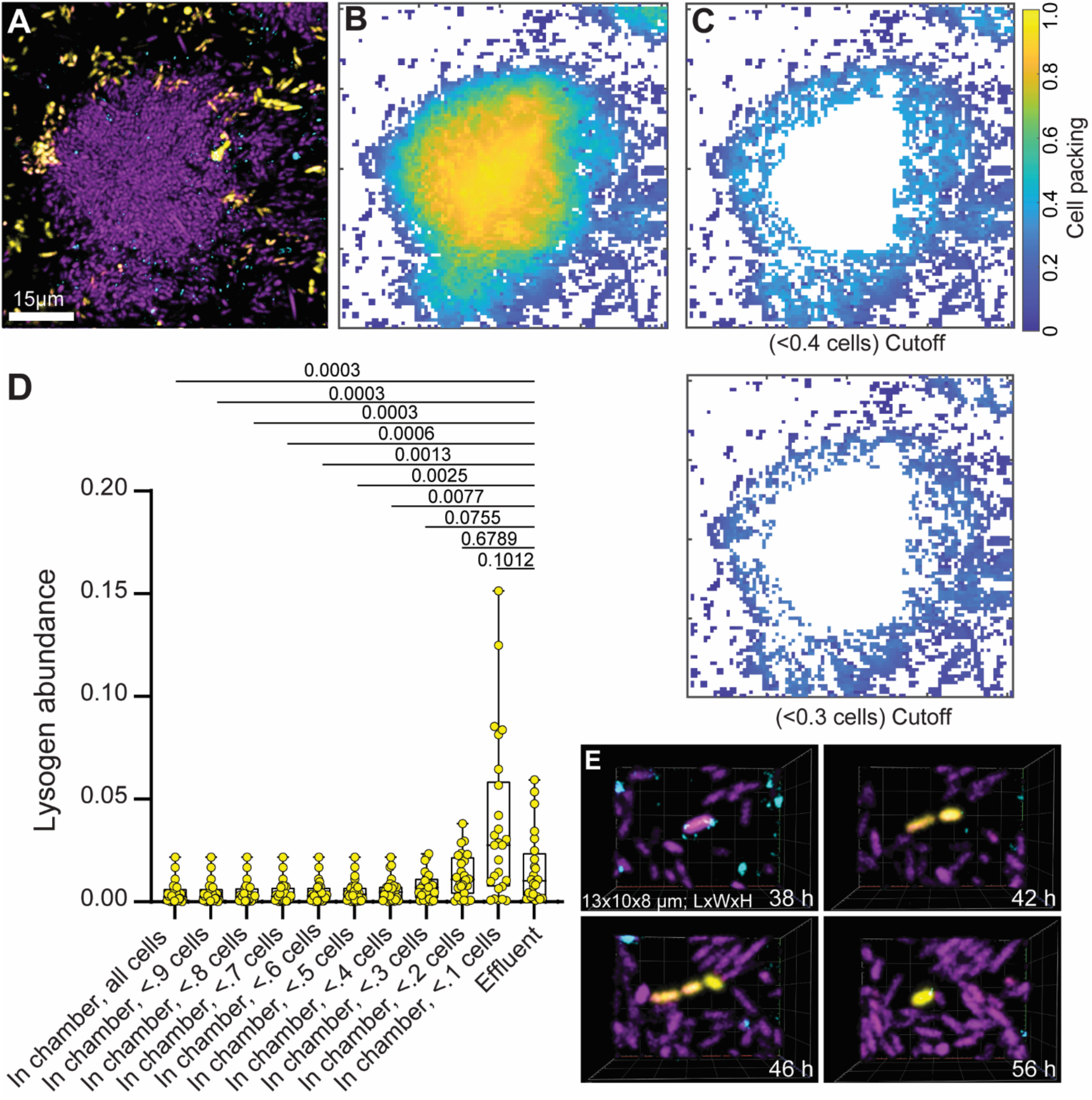
(A) Representative image of a biofilm chamber location when effluent was collected. (B) heatmap of cell packing density for the biofilm in panel (A). (C) Two Corresponding heatmaps of density of in panel (A), but with any cell volumes with cell packing >0.4 or >0.3 have been removed. (D) Lysogen abundance on a continuum of local densities compared to the effluent of the microfluidic device. Pairwise comparisons are Mann-Whitney U test with Bonferroni correction (E) Timelapse of a sensitive *E. coli* cell (purple) infected with λ (Turquoise) at 38 h and becoming lysogenized (yellow). We observed two division events before dispersal from the microfluidic device.

**SI Figure 14.**
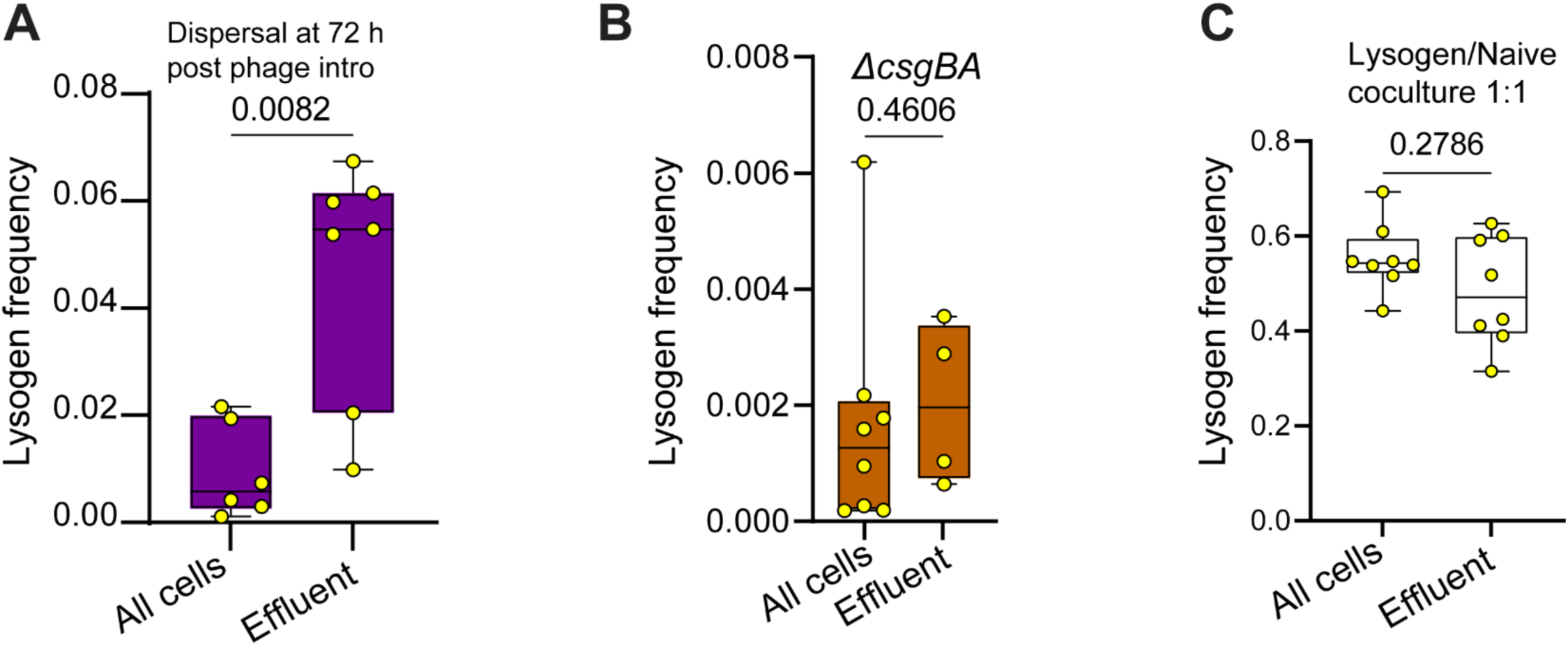
(A) Lysogen frequency of WT *E. coli* biofilms that have been invaded with λ phages for 72 h (n=6-7). (B) Lysogen frequency in Δ*csgBA* microfluidic device biofilms and dispersing effluent (n=4-8). (C) Lysogen frequency in the total biofilm population and in the dispersing effluent population from 72 h biofilms in which lysogens and naïve WT *E. coli* cells were inoculated at a 1:1 ratio from the beginning of biofilm growth (n=8). Statistical tests shown are Mann-Whitney U tests.

**SI Figure 15.**
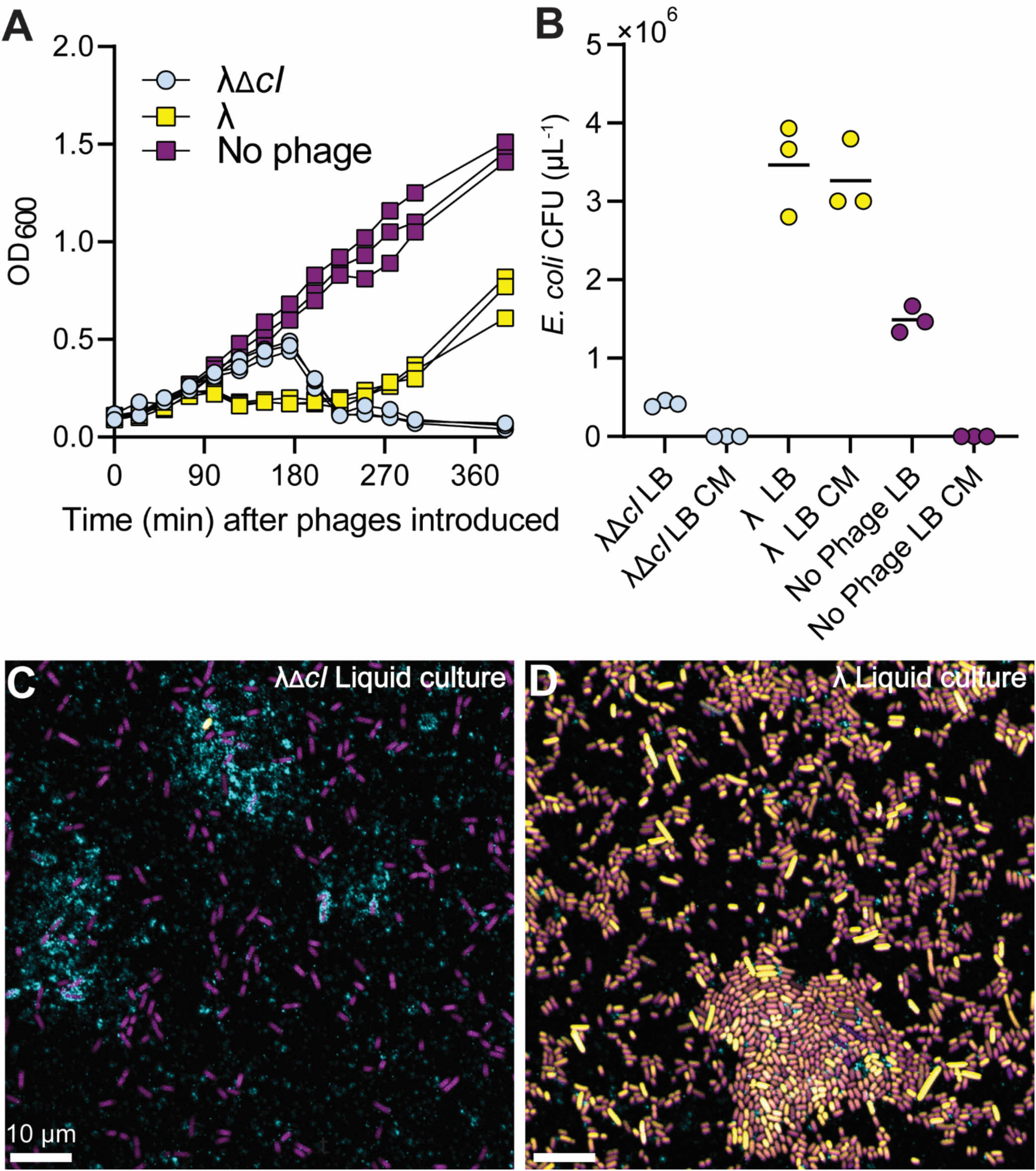
(A) OD600 curve of exponentially growing *E. coli* in λ broth, inoculated with λ or λΔ*cI* and grown at 30C°. After a lag period relative to the no-phage control, we observe an increase in OD600 of lysogens after addition of phage λ, but no increase in population density after addition of the obligately lytic phage derivative λΔ*cI*. (B) Colony Forming Units (CFUs) of *E. coli* on LB plates or LB + chloramphenicol (CM) plates after incubation with λ, λΔ*cI*, or no phage in LB media for 24 h, a separate experimental setup from that shown in panel (A). Our strain of phage λ contains a chloramphenicol resistance marker, so lysogens are Cm resistant. (C,D) Images of a sample taken from the λΔ*cI*/*E. coli* liquid culture and λ/*E. coli* liquid culture after 24 h.

## Notes

### Competing Interest Statement

The authors have declared no competing interest.

### Summary of Updates

- Updates and clarifications to main text and SI - Updates to figure composition; former figures 1 and 2 now combined into new figure 1 - Addition of new experiments and new figure 3 - Addition of new final supplemental figure

